# Field Modeling Study of Yield Response and Nitrate Leaching with Manure Application and Deficit Irrigation for Maize-Fallow-Wheat Rotation

**DOI:** 10.64898/2026.08.13.744693

**Authors:** Muhammad Tahir, David Mulla, Saliha Maqbool, Anwar Ul Hassan

## Abstract

Efficient nutrient and water management is crucial for enhancing crop productivity, soil health, and mitigating environmental losses in cereal cropping systems of semi-arid regions. A two-year field experiment was conducted to evaluate the effects of dairy manure annual application of 50 Mg ha^-1^ to maintain recommended fertilizer N, compared to sole urea application with two different irrigation regimes (100% and 75% ET_c_) on crop yield, water use efficiency, deep percolation, nitrate leaching, and soil quality within a wheat–fallow–maize rotation in Pakistan. Suction lysimeters installed at a depth of 1.2 m were used to collect nitrate-N leachates, while HYDRUS-1D was used to assess daily deep percolation losses. Results indicate that the interaction between manure and irrigation was significant for yield, nitrate-N leaching, and soil health. Manure with deficit irrigation showed wheat and maize yield of 4.36 and 7.80 Mg/ha, irrigation water use efficiency (WUE_i_) of 1.09 and 1.59 kg/ha/mm, respectively, with no significant increase observed with full irrigation; while a significant decrease was observed in the absence of manure, either with full irrigation or deficit irrigation. Manure with deficit irrigation averaged annual nitrate-N leaching of 17.45 kg/ha, while urea and manure with full irrigation averaged 11.46 and 55.59% increases in nitrate-N leaching losses, respectively, without any yield benefits. Our results indicate that deficit irrigation with manure produces optimum yield with reduced nitrate-N leaching risk and improved soil physical properties.

## 1. Introduction

Water-fertilizer coupling has emerged as a pivotal strategy in modern agriculture for its potential to enhance crop yield, reduce NO_3_-N leaching losses, improve soil health, and ensure better resource efficiency (Xing et al., 2024). Nitrogen (N) addition to row cropping systems is a vital facet of agriculture, where the increasing use of N fertilizers is a key factor in intensifying agricultural production and meeting global yield increase demands. However, high-production cropping systems cause substantial N leaching losses due to the difficulty in predicting site-specific crop N requirements and losses during the fallow period. Where high rates of N fertilizers are applied to maize, relatively low recovery efficiencies ranging from 35-60% have been observed (Randall et al., 1997; van Es et al., 2019). Moreover, varying weather conditions, especially high-intensity rainfall during the fallow period, result in high-water infiltration and nitrate-N leaching losses (Tahir et al., 2025). Globally, both wheat and maize exhibit low crop water productivity, presenting a significant opportunity to enhance agricultural production with reduced water consumption (Bastiaanssen and Steduto, 2017). Inappropriate N fertilizer and water management practices are causing yield instability in Pakistan. N fertilizer application in Pakistan’s agricultural system has increased more than tenfold during the past six decades to boost yield; however, this has resulted in environmental degradation. Intensive irrigation and fertilizer use in Pakistan are thus leading to groundwater pollution, particularly from N fertilizer (Leghari et al., 2024; Tahir et al., 2012a). Shallow groundwater pollution by nitrate is observed in Pakistan. In Punjab, flood irrigation and rainfall patterns exceed crop water uptake rates, causing leaching of water and solutes (Afzal et al., 2000).

Several factors contribute to the low yields of wheat and maize in Pakistan. Nitrogen fertilizer recovery rates are low, and flood irrigation, along with fallow periods in crop rotation, contributes to nitrate leaching. Agricultural soils in Pakistan typically have low organic carbon and nitrogen content, which ultimately limits crop productivity. Organic amendments like manure are thus gaining interest in improving soil fertility and crop production (Khan et al., 2007). Organic sources of N, such as manure, not only increase crop yield in low soil organic matter (SOM) but also improve the physicochemical properties of the soil. Manure application in semiarid conditions of Pakistan was observed to improve water- and fertilizer-use efficiency, improve soil properties, and increase economic return compared to NPK application. Moreover, residual soil organic fertility following crop harvest was equivalent to that obtained from fresh dairy manure application (Tahir et al., 2012a; Afzal et al., 2000; Khan et al., 2007; Azam, 1988; Tahir et al., 2012b).

Manure, though commonly used, faces constraints due to its use as fuel, impacting its availability. However, combining farm manure with inorganic fertilizers has been shown to increase crop yields while minimizing nitrate leaching (Tahir et al., 2012; Khan et al., 2007). Lack of efficient irrigation management is one of the major reasons for the low crop yield and negative environmental impacts of irrigated agriculture. Precise irrigation with optimum N fertilization not only increases yield but also reduces nitrate-N leaching risks (Muhammad et al., 2022; Arif et al., 2016; Singh et al., 2023). A relationship between NO_3_ - leaching and water application amounts using the HYDRUS model indicated that NO_3_ - leaching can be decreased by 20–48 % under a 10–20 % deficit water application (Khan et al., 2019). Nitrogen in the form of nitrate is highly mobile in soil and is primarily influenced by soil water conditions. High water is needed for better crop growth and development; however, it causes potential N losses in the form of N leaching. Leaching of nitrate-N below the crop root zone can be minimized by managing N and water inputs to the crop simultaneously (Muhammad et al., 2022). Thus, both N fertilization and water management are necessary to decrease potential nitrate losses without affecting maize growth and performance (Tahir et al., 2012).

Pakistan is one of the world’s most arid countries with an average yearly precipitation of approximately 250 mm, while it allocates up to 94% of its fresh water for agricultural use. Yet inefficient irrigation scheduling and outdated technologies hinder proper management, resulting in low water use efficiency (WUE) and crop yields (UNEP, 2011; FAO, 2016; Jabeen et al., 2022). Since high nitrogen application rates are essential in low SOM soils to achieve maximum crop yield, proper scheduling alongside irrigation events is essential to ensure plant access to nitrogen. Understanding soil water dynamics and implementing optimal deficit irrigation schedules are crucial for efficient water conservation (Eghball et al., 2004; Doorenbos and Kassam, 1979). Moreover, a significant amount of NO_3_-N leaching occurs during the fallow monsoon rainy season in Punjab, Pakistan. Studying NO_3_- leaching during fallow periods is crucial, with soil texture and organic amendments affecting leaching rates. Extensive in situ measurement of soil moisture is time-consuming and cost-prohibitive. Alternatively, different simulation models, when locally calibrated, can accurately assess the soil water balance. The HYDRUS-1D model has been used worldwide to simulate soil water dynamics and water fluxes under different land-use types. This model allows the specification of root water uptake, evaporation during the fallow period, and adjustment of soil hydraulic properties for different scenarios (Šimunek et al., 2011; Tahir et al., 2016).

The main objectives of the study were: I) Assessing the impact of immediate and residual effects of manure vs sole urea application under full and deficit irrigation on crop yield, soil physical properties, and nitrate leaching losses. II) Assessing the commonly used dose of 50 mg ha^-1^ fresh manure for crop N requirement, yield, and nitrate leaching. III) Evaluating the profitability of manure application under deficit irrigation conditions. IV) Quantifying the amount of nitrate leaching and drainage during the wheat, maize, and fallow periods.

## 2. Materials and Methods

### 2.1. Experimental site and climatic conditions

The field experiments were conducted for two years (2007-08 to 2008-09), using wheat-fallow-maize rotation at Research Farm (31°-26’ N and 73°-06’ E; altitude, 184.4 m), Institute of Soil and Environmental Sciences, University of Agriculture, Faisalabad, Punjab, Pakistan. The climate of the area is sub-tropical. The annual mean temperature is 31.6°C, with maximum and minimum monthly average temperatures of 41°C and 19.4°C observed during the month of June and January, respectively. The average annual precipitation of this area is about 684 mm. During the study period of 2007-08 and 2008-09, the experimental site received 517.1 mm and 373.4 mm precipitation, respectively, with 71.8% and 61.2% falling only during the fallow period (Figure 1). The crop was thus dependent on irrigation. The soil of the experimental site (Table S1) is well-drained Hafizabad loam, mixed, semi-active, isohyperthermic Typic Calciargids. Soil is low in soil organic carbon (SOC), alkaline in nature (pH 8), and contains medium soil water retention capacity. Soil properties were measured at depths of 0.0-0.35 m, 0.35-0.70 m, and 0.70-1.2 m (Table 1). The sand proportion of the soil slightly increased for lower depths; however, the soil texture remained the same, i.e., loam. Soil bulk density of the surface 0-0.35 m depth was 1.5 Mg m-3, with a slight increase observed for lower depths. Soil Water contents at saturation, field capacity, and permanent wilting point for different depths were observed in the range of 0.43-0.42 m3 m-3, 0.265-0.254 m3 m-3, and 0.123-0.126 m^3^ m^-3^, respectively. Soil saturated hydraulic conductivity was higher (30.4 cm day^-1^) near the surface and decreased with depth. SOC was observed at 0.67 and 0.35% for 0-0.35 m soil with and without manure amendment, respectively. Lower depths of 0.35-0.70 and 0.7-1.2 m showed decreased values of 0.28 and 0.22%, respectively.

**Figure 1.**
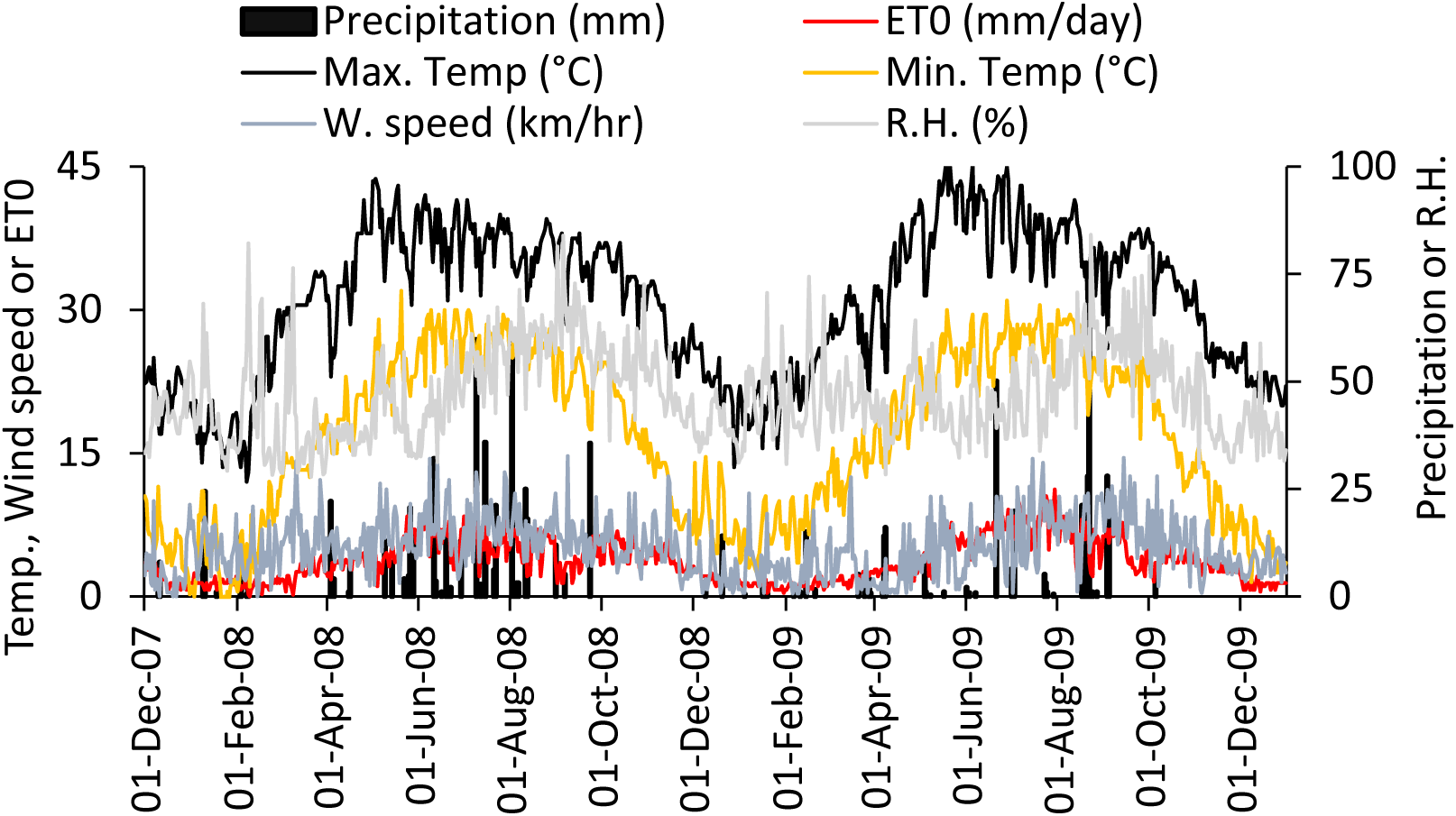
Precipitation and climatic conditions during the study period.

**Table 1.** Measured soil physical and hydraulic parameters in the three main layers of the experimental site.

| Depth<br>cm | Particle fraction (%) | | | B.D.<br>(Mg m<sup>-3</sup>) | $\theta_s$<br>-----cm<sup>3</sup> cm<sup>-3</sup>----- | $\theta_{FC}$ | $\theta_{PWP}$ | $K_s$<br>cm d<sup>-1</sup> | SOC<br>(%) | |
| --- | --- | --- | --- | --- | --- | --- | --- | --- | --- | --- |
| 0-35* | 38.0± | 37.5± | 24.5± | 1.50± | 0.43± | 0.265± | 0.123± | 30.4± | 0.67± |  |
|  | 0.13¶ | 0.14 | 0.10 | 0.03 | 0.02 | 0.008 | 0.015 | 0.45 | 0.10 |  |
| 0-35 | 38.0± | 37.0± | 25.0± | 1.51± | 0.43± | 0.260± | 0.125± | 27.3± | 0.35± |  |
|  | 0.09 | 0.13 | 0.15 | 0.02 | 0.01 | 0.010 | 0.013 | 0.52 | 0.03 |  |
| 35-70 | 40.0± | 34.5± | 25.5± | 1.55± | 0.42± | 0.256± | 0.127± | 19.5± | 0.28± |  |
|  | 0.21 | 0.25 | 0.13 | 0.01 | 0.01 | 0.013 | 0.014 | 0.45 | 0.02 |  |
| 70-110 | 42.5± | 32.5± | 25.0± | 1.54± | 0.42± | 0.254± | 0.126± | 20.0± | 0.22± | * Texture was loam according to |
|  | 0.15 | 0.16 | 0.15 | 0.02 | 0.01 | 0.012 | 0.009 | 0.40 | 0.01 | USDA, ¶Mean± standard error (data |

are average of three repeats); B.D., bulk density; $\theta_s$ , soil water content at saturation; $\theta_{FC}$ , soil water content at field capacity; $\theta_{PWP}$ , soil water content at permanent wilting point; $K_{fs}$ , soil saturated hydraulic conductivity.

### 2.2. Dairy manure and irrigation treatments

Fresh dairy manure was collected from dairies located around the University of Agriculture, Faisalabad, Pakistan. At the time of field application, fresh dairy manure samples were collected and later analyzed for organic carbon and nutrient (nitrogen, Phosphorus, and potash) contents, and soil moisture contents were measured using methods described by Nelson and Sommers (Nelson and Sommers, 1982). Results indicated that there was only a slight difference in organic carbon and nutrient contents of dairy manure (Table S2). During 2008 and 2009, dairy manure contained 60.5 and 68.4% moisture, 1.42 and 1.38% nitrogen (N), 0.47 and 0.50% phosphorus (P) as P_2_O_5_, and 1.3 and 1.2% potash (K) as K_2_O. Two rates of manure (0 and 50 Mg ha^-1^) were applied to wheat, while the residual effect of manure was assessed on the maize crop. The availability of N contents from fresh dairy manure was considered 55% for the first crop (wheat). It was estimated that 276 and 227 kg N, 94 and 80 kg P_2_O_5_, and 240 and 208 kg K_2_O were present in manure during 2007-8 and 2008-9, while it was assumed that only 55% was available for the first crop (wheat). The N rate to the wheat crop was adjusted based on the N contents in manure and percent availability. The manure was incorporated at 0-15 cm during land preparation of wheat in both years, and its residual effect was observed on the maize crop in both years. No adjustment was made to N rates of maize crops. For the wheat crop, a recommended dose of NPK was maintained at the rate of 105-85-62 kg N-P_2_O_5_-K_2_O ha^-1^. Wheat variety AS-2002 was used as the test crop, planted at a 9-inch row-to-row distance using a hand drill. A recommended dose of NPK was applied to the maize crop at 195-140-105 kg N-P_2_O_5_-K_2_O ha^-1^. Hybrid maize variety Pioneer-3062 was sown as the test crop, planted at 22 cm plant-to-plant and 67 cm row-to-row distance (36 plants per lysimeter). Maize did not receive any manure while the residual effect of M50 was studied. The wheat and maize trials were repeated the next year, with the same layout.

The two irrigation levels were 325 mm and 475 mm for wheat, and 450 and 600 mm for maize, corresponding to 75%ETc and 100%ETc, respectively. In addition, the experimental site received 75.4, 40.5, 80.3, and 113.8 mm precipitation during wheat 2007-8, maize-2008 wheat-2008-9, and maize-2009 crop, respectively. All irrigations in the field were maintained with a cut-throat flume. The Fallow period received 371 and 228 mm precipitation in addition to one irrigation of 100 mm for the purpose of sowing.

### 2.3. Dairy manure and irrigation treatments

The soil was tilled to a depth of 30 cm. Plots measuring 6.7 m x 13.3 m were prepared. The manure and irrigation were applied according to a split-plot design, where manure levels were applied in main plots, while the irrigation levels were applied in subplots. Wheat and maize field trials (Table S2) were conducted in plots of 6.7 m x 13.3 m dimensions, with the same treatment following a split-plot design, using two manure levels (0 and 50 t ha^-1^) in main plots and two irrigation levels in subplots, with six replicates. The first-year wheat crop (wheat-yr1) was sown on Dec. 1, 2007, and harvested on Apr. 19, 2008 (141 days), while during the 2nd year wheat (wheat-yr2) was sown on Dec. 16, 2008, and harvested on Apr. 27, 2009 (133 days). There was a fallow period of 122 and 125 days following the wheat crop during 2008 and 2009, respectively. The first-year maize (maize-yr1) crop was sown on Aug. 20, 2008, and harvested on Dec.12, 2008 (115 days), while 2nd year maize crop (maize-yr2) was sown on Sept. 1, 2009, and harvested on Dec. 23, 2009 (112 days). Isoproturon 50 WP was applied to the wheat crop using a knapsack sprayer at 55 days after sowing (DAS), following the second irrigation, at a rate of 1.0 kg active ingredient (a.i.) ha^-1^. Atrazine 38 SC was applied to the maize crop at 30 DAS at a rate of 0.774 kg a.i. ha^-1^.

### 2.4. Data Collection

#### 2.4.1. Plant measurement

The leaf area index (LAI) was measured by a digital leaf area meter. The leaf area index of the wheat crop was measured at 20, 30, 40, 50, 60, 70, 80,100, 110, and 120 days after sowing (DAS). Maize crop Leaf area index was measured at 10, 20, 30, 40, 50, 60, 70, 80, and 100 DAS, and at harvest. Plant height of wheat crop was measured using a meter tape at 20, 30, 50, 70 and 90 DAS, and at the time of harvest. Maize crop height was measured at 20, 30, 40, 50, 60, and 70 DAS, and at harvest. Root weight density (RWD) and Root length density (RLD) were measured for both wheat and maize crops.

Roots of both crops were sampled for RWD and RLD at the end of the mid-crop growth stage. Wheat root samples were taken within rows using a root sampler with a 15 cm depth and a 4 cm radius. Maize plant roots were sampled manually from 0.35 m^3^ by taking out a bulk of soil with the help of sampling tools. Roots were washed by root washing systems or manually, depending on feasibility. They were then dried at 65 °C till no change in weight (mostly for 24 hours) and then weighed (Rosario et al., 2000). Root length density of samples was measured with the help of Dalta T-Scan. For large maize samples, RLD was determined on a 5 g sub-sample. Wheat and maize crops were harvested at maturity from one square meter, and the yield of both crops was converted to Mg ha^-1^.

Water-use efficiency (WUE) was measured as (Hussain et al., 1995):

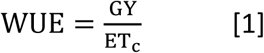

Where WUE (kg ha^-1^ mm^-1^) is the water use efficiency for grain yield (kg ha^-1^), GY is the grain yield (kg), and ETc (mm), which was calculated by the Panman-Montieth FAO-56 Method. Irrigation water use efficiency (WUE_i_) was calculated as follows:

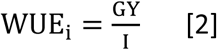

#### 2.4.2. Leachate collection, NO_3_-N measurement and drainage calculation

In field trials in which crops were grown, ceramic cups were inserted at three depths, i.e., 0.35, 0.70 and 1.1 m in each plot (Figure S1). These ceramic cups were buried after the sowing of the 1st wheat crop. These were installed randomly in the plots, at least 2 m away from the edges. Ceramic cups were prepared and inserted according to the method stated by (Webster et al., 1993). All the samplers were pre-treated using 1 M HCl as stated by (Debyle et al., 1988). Leachates were collected at predefined intervals (Table S3) by applying 0.6 to 0.7 bar suction with the help of a suction pump (Fig. 3.3). Samples were analyzed within 24 hrs of leachate collection, otherwise stored at −4 °C. Nitrate leaching at different depths was calculated based on the following equations:

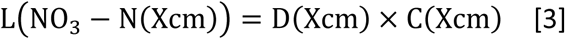

Where D is drainage, and C is nitrate concentration in leachates, while x cm is the depth in cm, i.e., 70 and 110 cm. To measure nitrate build-up, soil samples from three depths, 0-35 (D1), 35-70 (D2), and 70-110 cm (D3), were collected and analyzed for NO_3_-N contents. The NO_3_-N at predefined time intervals was calculated by subtracting the current NO_3_-N mass from the initial budget. The NO_3_-N mass (kg ha^-1^) up to a certain depth of soil was calculated by multiplying NO_3_-N concentration (mg kg^-1^) by the soil mass of one hectare for respective depths. Nitrate-N was analyzed by the chromotropic acid method (Hadjidemetriou, 1982). To measure soil NO_3_-N, 10 g of an air-dried sample was placed in an Erlenmeyer flask, and 50 mL of 0.01 N copper sulfate solution was added. The mixture was shaken for 15 minutes and then filtered through Whatman No. 42 filter paper. Then, 3 mL of the filtrate was taken in a 10 mL volumetric flask, to which 1 mL of 0.1% chromotropic acid and 6 mL of concentrated sulfuric acid were added. The NO_3_-N concentration was then determined at 430 nm using a spectrophotometer. Leachate NO_3_-N concentration (ppm) was measured by taking a 1 mL leachate sample, while the rest of the procedure remained the same as for the soil samples. An initial estimate of drainage was made before each irrigation event, using the following water balance equation:

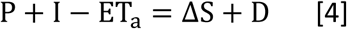

Here, I is the depth of irrigation applied (mm), which was calculated using a cut throat flume; ET_a_ is the actual evapotranspiration (mm day^-1^); P is the precipitation (mm; ΔS is the change in water storage (mm) of the soil profile up to the required depth. Water contents were measured gravimetrically by taking soil samples at each depth before each irrigation, before the start and at the end of the trial; D is the drainage or deep percolation (mm). Actual evapotranspiration was calculated by the Penman-Monteith FAO-56 Method (Allen et al., 1998).

For the measurement of NO_3_-N leaching losses on a daily basis, daily deep drainage was simulated using the HYDRUS-1D model. HYDRUS-1D was calibrated using the soil properties and soil water retention parameters of different soil layers (Table 1). The dual-porosity model was used to describe water flow, which is based on the Richards equation (Simunek et al., 2003), where a mass balance equation was used to illustrate moisture dynamics in soil as follows:

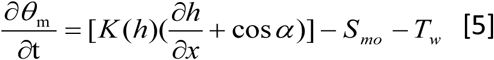

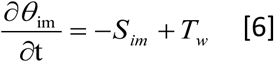

where S_m_ and S_im_ are root water uptake for these two regions, and T_w_ is the water transfer rate from the mobile to immobile region (inter to intra-aggregate pores), and 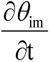 is the change in immobile water contents with respect to time.

#### 2.4.3. Soil properties measurement

Oxidizable soil organic carbon (SOC) was analyzed using the standard procedure given by (Ryan et al., 2001). Soil bulk density from 0-5, 5-10, 10-20, and 20-30 cm depths was determined by the core sampler method as described by (Blake and Hartage, 1986). The percentage of sand, silt, and clay was measured by the Bouyoucos hydrometer method, and textural class was determined using the International Textural Triangle (Ryan et al., 2001).

Infiltration rate was measured with a double ring infiltrometer (Figure S1). The inner and outer rings were driven about 10 cm into the soil using an im-pact-absorbing hammer and a driving plate, and then they were filled with water. The water flowing through the inner ring into the soil was noted until a constant rate was achieved (Blake and Hartage, 1986). Soil saturated hydraulic conductivity (K_fs_) was measured (Figure S1) by a Guelph Permeameter (Model 2800 KI), taking three steady-state readings. The K_fs_ was then calculated from the following formula:

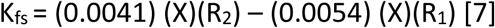

Where R_1_ and R_2_ are the steady-state rates of water flow (cm s^-1^) in the reservoir at the first (h_1_) and second head (h_2_) of water (cm), respectively, and X (35.5 cm^2^) is the reservoir constant, which is related to the cross-sectional area of the combined reservoir (cm^2^).

#### 2.4.4. Measurement of soil water retention parameters

Water retention curve (van Genuchten parameters) was measured for soil layers of 0-35, 35-70, 70-110 cm from field trials, while from 0-35, 35-70, 70-115 and 115-160 cm from lysimeters (Table 1). From manure-receiving lysimeter plots and field trials, soil samples were collected separately at 0-35 cm to find any change in water retention. Soil retention capacity was measured by determining water contents at pre-defined matric potential (Dane and Hopmans, 2002) with the help of suction plates at 0.3, 0.6, 1.0, 3.0, and 4.5 bar pressure. The recorded data were fitted to the dual-porosity model (Durner, 1994) using RETC-fit version 6.02 software to determine hydraulic parameters in the three main layers of the experimental site.

### 2.5. Statistical Analysis

The data collected were statistically analyzed using ANOVA (analysis of variance) techniques according to RCBD with split plot arrangement. The means were compared by LSD (least significant difference) test at p≤ 0.05. (Steel et al., 1997). The software packages STATISTIX 8.1 (StatSoft, Inc., 2001) and STATISTICA (Version 8, www.statsoft.com, OK 74104, US) were used for statistical analysis. Data was analyzed for year effect using two-tailed Student distribution (t-test).

## 3. Results

### 3.1. Grain yield, irrigation water use efficiency, and water use efficiency

The individual effects of manure and irrigation were statistically nonsignificant, while the interactive effect of manure and irrigation on grain yield and irrigation water use efficiency (WUE_i_) was statistically significant (p<0.05) (Table 2). Manure application increased grain yield and water use efficiency compared to no manure application in both growing seasons of wheat and maize. Grain yield increased with full irrigation (ET100%) compared to deficit irrigation (ET75%); however, it resulted in a lower WUE_i_. The highest grain yields in both wheat and maize growing seasons were obtained with the combined application of manure and full irrigation. However, these yields were not significantly different from those achieved with manure application under deficit irrigation, except during the first year of wheat (Wheat-Yr1). Application of manure at deficit irrigation, averaged over two years, resulted in wheat and maize yields of 4.36 and 7.80 Mg ha^-1,^ with no significant increase observed with 75% ET_c_. Conversely, Full irrigation and deficit irrigation without manure showed a 20.0 and 31.9% decrease in wheat yield, while 13.3 and 23.5% decreases in maize yield, respectively. Moreover, manure (M50) at deficit irrigation significantly increased the WUE of wheat by 18.9 and 20.2%; and maize by 5.6 and 28.8%, depicting values of 0.96 and 1.85 kg ha^-1^ mm^-1^, compared to those observed with urea application (M0) at full and deficit irrigation, respectively (Table S4). Full irrigation and deficit irrigation showed statistically similar WUE in the presence of manure. Significantly higher WUE_i_ of 1.09 and 1.59 for wheat and maize were observed with deficit irrigation in the presence of manure, respectively. Comparatively, manures with full irrigation to wheat and corn showed a decrease of 22.9 and 19.5%; urea with deficit irrigation showed a decrease of 32.1 and 23.6%; urea with full irrigation showed a decrease of 42.2 and 33.7%. Moisture measured (Figure S2) before each irrigation, and observed critical limit of readily available water (θ_RAW_) indicates that deficit irrigation caused water to drop below the θ_RAW_ leading to soil water stress to plant.

**Table 2.** Grain yield and irrigation water use efficiency of wheat and maize crops during different years under different irrigation and manure treatments.

|  | Grain yield (Mg ha<sup>-1</sup>) |  |  |  | Water use efficiency (kg ha<sup>-1</sup> mm<sup>-1</sup>) |  |  |  |
| --- | --- | --- | --- | --- | --- | --- | --- | --- |
|  | Wheat-1 | Wheat-2 | Maize-1 | Maize-2 | Wheat-1 | Wheat-2 | Maize-1 | Maize-2 |
| M<sub>0</sub>* | 3.36 | 3.09 | 6.25 | 6.48 | 0.72 | 0.66 | 1.11 | 1.16 |
| M<sub>50</sub> | 4.56 | 4.42 | 8.12 | 7.88 | 0.98 | 0.95 | 1.46 | 1.41 |
| I<sub>1</sub>** | 3.86 | 3.91 | 6.42 | 6.22 | 0.97 | 0.98 | 1.31 | 1.27 |
| I<sub>2</sub> | 4.41 | 4.37 | 6.72 | 6.6 | 0.8 | 0.8 | 1.05 | 1.03 |
| M<sub>0</sub>I<sub>1</sub> | 3.17 c° | 2.76 d | 5.81 b | 6.12 c | 0.79 b | 0.69 c | 1.18 bc | 1.25 bc |
| M<sub>0</sub>I<sub>2</sub> | 3.55 b | 3.42 c | 6.69 b | 6.83 bc | 0.64 c | 0.62 d | 1.04 c | 1.07 c |
| M<sub>50</sub>I<sub>1</sub> | 4.44 a | 4.27 b | 8.05 a | 7.55 ab | 1.11 a | 1.07 a | 1.64 a | 1.54 a |
| M<sub>50</sub>I<sub>2</sub> | 4.67 a | 4.56 a | 8.19 a | 8.20 a | 0.85 b | 0.83 b | 1.28 b | 1.28 b |
| LSD M | 0.34 | 0.43 | 0.89 | 0.74 | 0.08 | 0.1 | 0.16 | 0.13 |
| I | 0.2 | 0.16 | NS | NS | 0.05 | 0.04 | 0.17 | 0.16 |
| M x I | 0.28,0.39¶ | 0.23,0.46 | 1.38,1.32 | 1.23,1.14 | 0.07,0.09 | 0.06,0.1 | 0.25,0.23 | 0.23,0.20 |

1st LSD
0.05) value is for same levels of Manure, while 2nd for different levels of manure; Statistically (T-test) similar grain yield and WUE of wheat and maize was observed during both years

### 3.2. Crop evapotranspiration and fallow period evaporation

The main effects of manure and irrigation were statistically non-significant, while the interactive effect of manure and irrigation was statistically (p < 0.05) significant on crop evapotranspiration (ET_c_) during both the growing seasons of maize and wheat. No significant treatment effect was observed on evaporation during the fallow period (Table 3). Overall, applying manure to the wheat crop and its residual effect on the maize crop under full irrigation resulted in higher evapotranspiration compared to no manure application under deficit irrigation. Full irrigation with manure application showed the highest ET_c_ of 505-, 485-, 476-, and 445-mm during wheat-Yr1, maize-Yr1, wheat-Yr2, and maize-Yr2, respectively. However, these values were not significantly different from those obtained under reduced irrigation with manure application, except in the second year of maize (Maize-Yr2). ET_c_ observed under deficit irrigation with manure application was comparable to full irrigation under no manure. Full irrigation with manure application showed the highest increase of 29.8, 16.9, 20.8, and 21.6% for wheat-Yr1, maize-Yr1, wheat-Yr2, and maize-Yr2, respectively, compared to deficit irrigation without any manure application. Evaporation during the fallow period in Yr1 and Yr2 contributed 21.7 and 14.8% of the annual ET_c_, respectively. The main effect of manure and irrigation was non-significant, while the interaction effect of manure × irrigation was significant on drainage at p < 0.05 (Table 4) in both years of wheat and maize cultivation at depths of 70 cm and 110 cm. Manure application decreased drainage in both wheat and maize growing seasons. Drainage increased with deficit irrigation in both wheat and maize cultivation years and at various soil depths. Overall, the highest drainage was recorded in plots where no manure with deficit irrigation was applied across both crop seasons. Drainage decreased with depth, with higher drainage recorded at 70 cm soil depth compared to 110 cm depth. At 70 cm soil depth, compared to no manure under full irrigation, the application of manure under deficit irrigation showed an increase in drainage of 94.9 and 68.4% in Wheat-Yr1 and Wheat-Yr2, respectively. Meanwhile, the residual effect of manure with deficit irrigation resulted in an increase in drainage by 61.8 and 10.0% in Maize-Yr1 and Maize-Yr2, respectively (Table 4).

**Table 3.** Effect of manure and irrigation levels on evapotranspiration/ evaporation (cm) during wheat-fallow-maize crop rotation.

|  | Year-1 |  |  | Year-2 |  |  |
| --- | --- | --- | --- | --- | --- | --- |
|  | Wheat | Fallow¶ | Maize | Wheat | Fallow¶ | Maize |
| M<sub>0</sub>* | 421 | 243 | 429 | 420 | 144 | 384 |
| M<sub>50</sub> | 491 | 257 | 464 | 455 | 147 | 422 |
| I<sub>1</sub>** | 433 | 244 | 429 | 414 | 145 | 382 |
| I<sub>2</sub> | 479 | 256 | 464 | 461 | 146 | 424 |
| M<sub>0</sub>I<sub>1</sub> | 389 c° | 231 | 415 b | 394 b | 146 | 366 c |
| M<sub>0</sub>I<sub>2</sub> | 453 b | 254 | 443 ab | 445 a | 142 | 402 b |
| M<sub>50</sub>I<sub>1</sub> | 476 ab | 256 | 443 ab | 434 ab | 145 | 398 b |
| M<sub>50</sub>I<sub>2</sub> | 505 a | 257 | 485 a | 476 a | 149 | 445 a |
| LSD M | 3.4 | 4.3 | 8.9 | 7.4 | 0.8 | 1.0 |
| I | 2.0 | 1.6 | NS | NS | 0.5 | 0.4 |
| M x I | 2.8,3.9¶ | 2.3,4.6 | 13.8,13.2 | 12.3,11.4 | 0.7,0.9 | 0.6,1.0 |

1st LSD ( $p \leq 0.05$ ) value is for the same levels of Manure, while the 2nd is for different levels of manure; Year effect ( $T$ -test- $p \leq 0.05$ ) was statistically significant for fallow period-1 at all depths, and for maize-1 at 110 cm depth.

**Table 4.** Effect of manure and irrigation levels on drainage (mm) at 70 cm and 110 m depth during wheat-fallow-maize rotation Year effect (*T*-test-p≤ 0.05) was statistically significant for wheat, fallow period, and maize period.

|  | 70 cm |  |  |  |  |  | 110 cm |  |  |  |  |  |
| --- | --- | --- | --- | --- | --- | --- | --- | --- | --- | --- | --- | --- |
|  | Year-1 |  |  | Year-2 |  |  | Year-1 |  |  | Year-2 |  |  |
|  | wheat | Fallow | Maize | Wheat | Fallow | Maize | wheat | Fallow | Maize | Wheat | Fallow | Maize |
| $M_0^*$ | 137 | 119 | 125 | 155 | 102 | 160 | 114 | 72 | 88 | 130 | 55 | 115 |
| $M_{50}$ | 117 | 108 | 100 | 132 | 105 | 126 | 83 | 87 | 72 | 122 | 53 | 87 |
| $I_1^{**}$ | 88 | 113 | 87 | 105 | 103 | 88 | 64 | 82 | 56 | 91 | 55 | 68 |
| $I_2$ | 165 | 113 | 180 | 182 | 104 | 184 | 133 | 77 | 104 | 161 | 54 | 134 |
| $M_{0I_1}$ | 97 c | 114 | 98 c | 112 c | 105 | 89 | 84 c | 76 ab | 66 b | 94 b | 56 | 75 c |
| $M_{0I_2}$ | 176 a | 123 | 152 a | 198 a | 98 | 203 | 142 a | 67 b | 110 a | 165 a | 54 | 154 a |
| $M_{50I_1}$ | 79 d | 112 | 76 d | 98 c | 100 | 87 | 43 d | 87 a | 46 c | 87 b | 53 | 60 c |
| $M_{50I_2}$ | 154 b | 103 | 123 b | 165 b | 110 | 165 | 123 b | 87 a | 97 a | 156 a | 53 | 114 b |
| LSD M | 10.7 | NS | 15.6 | 173 | NS | 18.9 | 14.0 | 9.9 | 11 | 7.5 | NS | 16.1 |
| I | 11 | NS | 14.4 | 9.0 | NS | 13.9 | 12.2 | 9.4 | 7.6 | 12.7 | NS | 14.9 |
| $M \times I$ | 15.6,15.4 | NS | 20.4, 21.1 | 12.7,19.5 | NS | 19.7,23.4 | 17,18 | 13, 14 | 1.1,1.3 | 18,15 | NS | 21, 22 |

Note. Means sharing the same letter (s) do not differ significantly at $P < 0.05$ according to Least Significance Difference Test

### 3.3. Nitrate leaching and Nitrate build-up in soil

The main effect of manure and irrigation was statistically non-significant, while the interaction effect of manure × irrigation was significant on nitrate leaching in both wheat and maize cultivation at soil depths of 70 cm and 110 cm at p< 0.05 (Table 5). Manure application increased nitrate leaching by 37.5, 31.3, 43.6, and 9.3% in wheat yr1, maize yr1, wheat yr2, and maize yr2 compared to no manure application. Deficit irrigation resulted in a lower nitrate leaching rate compared to full irrigation. Higher drainage was recorded where manure was applied at deficit irrigation. Overall, higher nitrate leaching was recorded at a soil depth of 70 cm compared to 110 cm. At 70, compared to no manure under full irrigation, the application of manure under deficit irrigation showed an increase in nitrate leaching of 65.9% and 91.4% in wheat-Yr1 and wheat-Yr2, respectively. Meanwhile, the residual effect of manure with deficit irrigation resulted in an increase in nitrate leaching by 95.3% and 87.7% in maize-Yr1 and maize-Yr2, respectively (Table 5). At a soil depth of 70 cm, nitrate leaching was reduced in yr2 compared to yr1, but at 110 cm soil depth, nitrate leaching increased in Yr2 compared to Yr1. The main effect of manure and irrigation was statistically nonsignificant, while the interactive effect of manure and irrigation on nitrate buildup in both wheat and maize cultivation years was significant at p < 0.05 (Table 6). The application of manure increased nitrate buildup, while full irrigation decreased nitrate buildup. Higher nitrate buildup was recorded at the 60-day sampling compared to harvesting. During the follow-up period in both years and soil depths, the lowest buildup of nitrate was observed, except for manure with full irrigation and deficit irrigation in Yr1, where these plots showed higher nitrate buildup compared to the buildup at the harvest stage. The buildup of nitrate was lower in the wheat (2008-09) cultivation season compared to the wheat (2007-08) cultivation season. At 60 days, when compared to no manure under full irrigation, applying manure under deficit irrigation led to an 18.0% increase in nitrate buildup in wheat-Yr1 and a 19.3% increase in wheat-Yr2. Additionally, the lasting impact of manure under deficit irrigation resulted in a 2.9% increase in nitrate buildup in maize-Yr1 and a 9.3% increase in maize-Yr2.

**Table 5.** Effect of manure and irrigation levels on NO_3_-N leaching at 70 cm and 110 m depth during wheat-fallow-maize rotation.

|  | 70 cm |  |  |  |  |  | 110 cm |  |  |  |  |  |
| --- | --- | --- | --- | --- | --- | --- | --- | --- | --- | --- | --- | --- |
|  | Year-1 |  |  | Year-2 |  |  | Year-1 |  |  | Year-2 |  |  |
|  | wheat | Fallow | Maize | Wheat | Fallow | Maize | wheat | Fallow | Maize | Wheat | Fallow | Maize |
| M<sub>0</sub>* | 8.8 | 5.5 | 9.6 | 9.4 | 4.8 | 8.6 | 6.4 | 2.6 | 4.9 | 7.7 | 3.2 | 6.8 |
| M<sub>50</sub> | 12.1 | 11.2 | 12.6 | 13.5 | 9.6 | 9.4 | 7.3 | 6.1 | 6.8 | 11.8 | 6.1 | 6.5 |
| I<sub>1</sub>** | 8.2 | 8.1 | 8.4 | 8.6 | 7.1 | 6.5 | 4.4 | 4.3 | 5 | 7.1 | 4.9 | 4 |
| I<sub>2</sub> | 12.7 | 8.6 | 13.8 | 14.3 | 7.3 | 11.5 | 9.3 | 4.4 | 6.7 | 12.5 | 4.3 | 9.4 |
| M<sub>0</sub>I<sub>1</sub> | 7.3 c | 5.3 b | 8.2 c | 7.9 d | 4.4 b | 6.4 b | 4.8 c | 2.6 | 4.2 c | 5.8 c | 3.3 c | 3.8 c |
| M<sub>0</sub>I<sub>2</sub> | 10.3 b | 5.7 b | 10.9b | 10.8 b | 5.2 b | 10.8 a | 8.1 b | 2.5 | 5.6 b | 9.7 b | 3.1 c | 9.9 a |
| M<sub>50</sub>I<sub>1</sub> | 9.1 bc | 10.9 a | 8.5 c | 9.3 c | 9.8 a | 6.5 b | 4.0 c | 5.9 | 5.8 b | 8.4 b | 6.6 a | 4.2 c |
| M<sub>50</sub>I<sub>2</sub> | 15.1 a | 11.5 a | 16.6 a | 17.8 a | 9.3 a | 12.2 a | 10.5 a | 6.3 | 7.8 a | 15.3 a | 5.5 b | 8.9 b |
| LSD M | 1.09 | 1.81 | 2.13 | 0.67 | 0.64 | 1.64 | 1.55 | 0.43 | 0.95 | 1.28 | 0.77 | 0.71 |
| I | 1.48 | 0.8 | 0.95 | 1.15 | 1.32 | 1.31 | 1.04 | 0.41 | 0.44 | 1.1 | 0.42 | 0.7 |
| MxI | 2.10,<br>1.84 | 1.14,<br>1.98 | 1.35,<br>2.33 | 1.63,<br>1.33 | 1.87,<br>1.46 | 1.85,<br>2.09 | 1.47,<br>1.86 | 0.58,<br>0.59 | 0.62,<br>1.04 | 1.56,<br>1.68 | 0.59,<br>0.88 | 0.99,<br>0.10 |

Year effect ( $T$ -test- $p \leq 0.05$ ) was statistically significant for wheat, fallow period, and for maize; \*Means sharing the same letter
(s) do not differ significantly at $P < 0.05$ according to Least Significance Difference Test

**Table 6.**
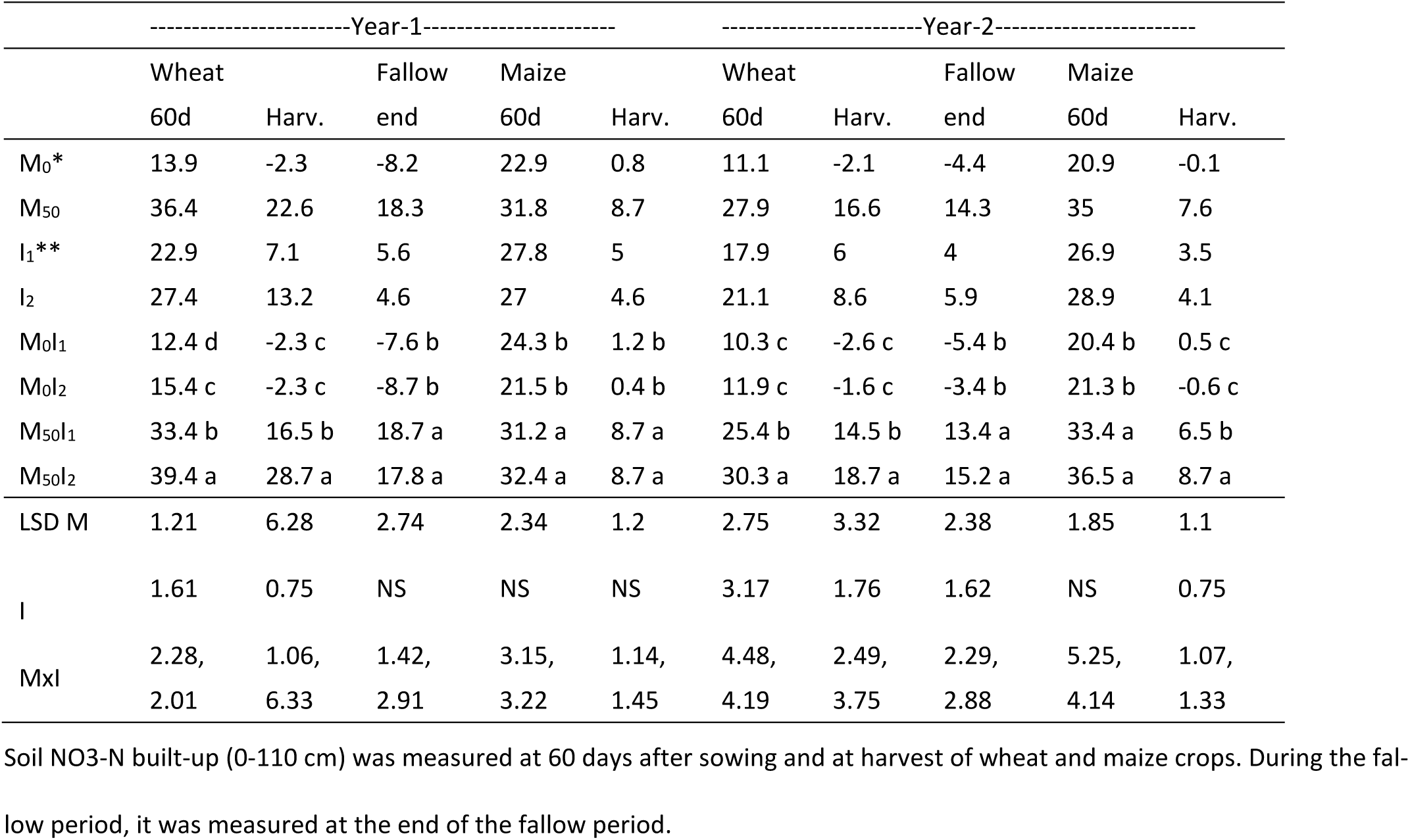
Effect of manure and irrigation levels on NO_3_-N built-up (kg ha^-1^) during wheat, fallow period and maize at 0-110 cm depths.

|  | -----Year-1----- |  |  |  |  | -----Year-2----- |  |  |  |  |
| --- | --- | --- | --- | --- | --- | --- | --- | --- | --- | --- |
|  | Wheat |  | Fallow | Maize |  | Wheat |  | Fallow | Maize |  |
|  | 60d | Harv. | end | 60d | Harv. | 60d | Harv. | end | 60d | Harv. |
| M<sub>0</sub>* | 13.9 | -2.3 | -8.2 | 22.9 | 0.8 | 11.1 | -2.1 | -4.4 | 20.9 | -0.1 |
| M<sub>50</sub> | 36.4 | 22.6 | 18.3 | 31.8 | 8.7 | 27.9 | 16.6 | 14.3 | 35 | 7.6 |
| I<sub>1</sub>** | 22.9 | 7.1 | 5.6 | 27.8 | 5 | 17.9 | 6 | 4 | 26.9 | 3.5 |
| I<sub>2</sub> | 27.4 | 13.2 | 4.6 | 27 | 4.6 | 21.1 | 8.6 | 5.9 | 28.9 | 4.1 |
| M<sub>0</sub>I<sub>1</sub> | 12.4 d | -2.3 c | -7.6 b | 24.3 b | 1.2 b | 10.3 c | -2.6 c | -5.4 b | 20.4 b | 0.5 c |
| M<sub>0</sub>I<sub>2</sub> | 15.4 c | -2.3 c | -8.7 b | 21.5 b | 0.4 b | 11.9 c | -1.6 c | -3.4 b | 21.3 b | -0.6 c |
| M<sub>50</sub>I<sub>1</sub> | 33.4 b | 16.5 b | 18.7 a | 31.2 a | 8.7 a | 25.4 b | 14.5 b | 13.4 a | 33.4 a | 6.5 b |
| M<sub>50</sub>I<sub>2</sub> | 39.4 a | 28.7 a | 17.8 a | 32.4 a | 8.7 a | 30.3 a | 18.7 a | 15.2 a | 36.5 a | 8.7 a |
| LSD M | 1.21 | 6.28 | 2.74 | 2.34 | 1.2 | 2.75 | 3.32 | 2.38 | 1.85 | 1.1 |
| I | 1.61 | 0.75 | NS | NS | NS | 3.17 | 1.76 | 1.62 | NS | 0.75 |
| MxI | 2.28, | 1.06, | 1.42, | 3.15, | 1.14, | 4.48, | 2.49, | 2.29, | 5.25, | 1.07, |
|  | 2.01 | 6.33 | 2.91 | 3.22 | 1.45 | 4.19 | 3.75 | 2.88 | 4.14 | 1.33 |

Soil NO<sub>3</sub>-N built-up (0-110 cm) was measured at 60 days after sowing and at harvest of wheat and maize crops. During the fallow period, it was measured at the end of the fallow period.

### 3.4. Root weight and root length density, Plant height, and leaf area index

The main effect of manure and irrigation was statistically non-significant, while the interactive effect of manure and irrigation was statistically significant at p<0.05 (Figure 4; Table S5). In both years of maize and wheat cultivation, manure with deficit irrigation led to an elevation in plant height compared to other treatments throughout all stages of height measurement, spanning from 20 days to the final plant assessment (Figure 1). On day 106, compared to no manure under full irrigation, the application of manure under deficit irrigation showed an increase in plant height of 1.3 and 0.9% in maize-Yr1 and maize-Yr2, respectively. Meanwhile, the residual effect of manure with deficit irrigation resulted in an increase in nitrate leaching by 3.8% in wheat Yr1. The individual impacts of manure application and irrigation did not yield statistically significant differences in leaf area index. However, their combined effect was statistically significant at the p < 0.05 level (Figure 2).

**Figure 2.**
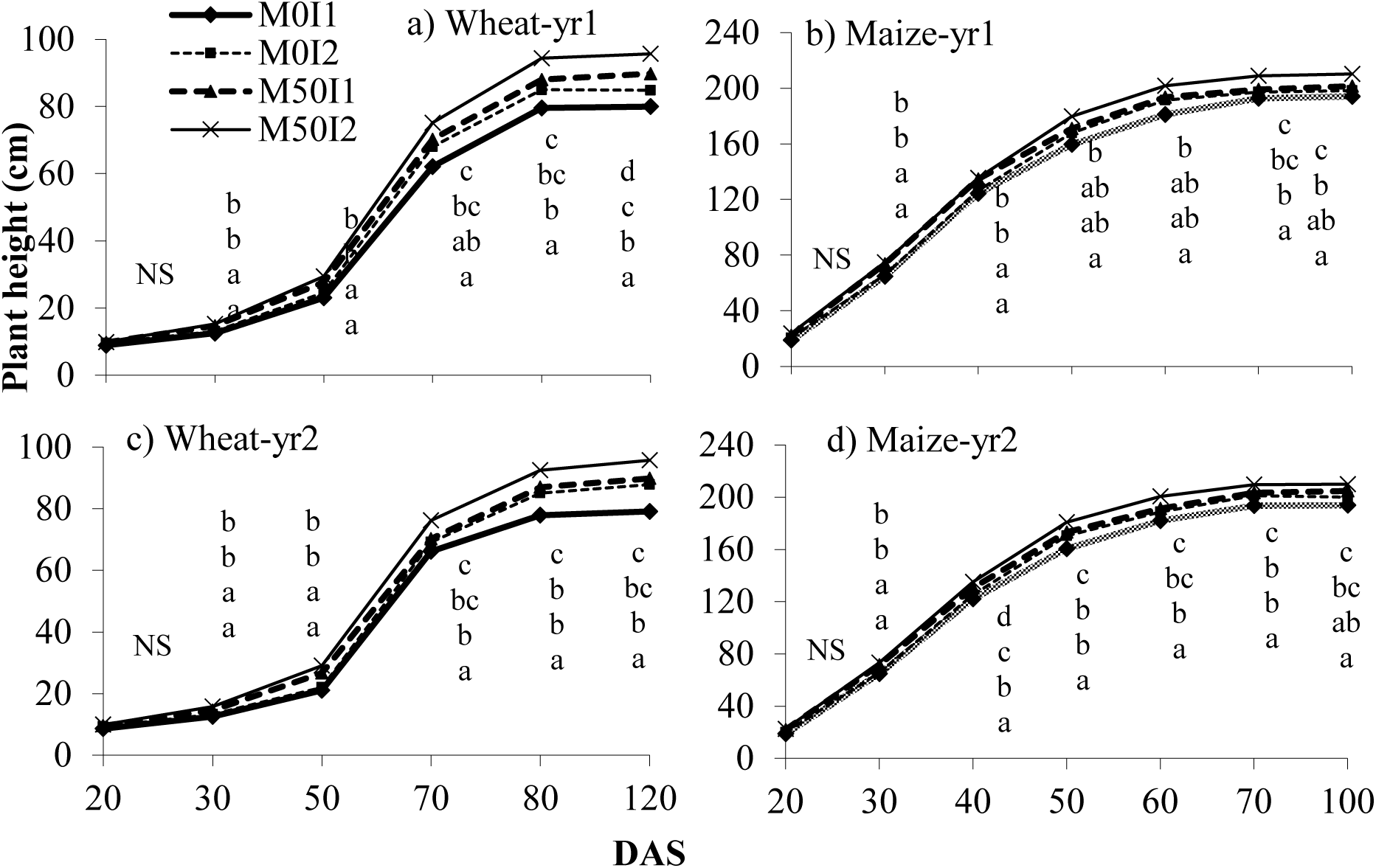
Effect of manure and irrigation on plant height measured at different time intervals during crop season Note. Year effect was statistically significant (*T*-test) for plant height at 40 days of yr-1 maize; and for LAI at 10 SAD of yr1 wheat, 30 DAS of yr-2 wheat and 70 DAS of yr-1 maize.

Manure application increased the leaf area index compared to scenarios without manure. Additionally, a higher leaf area index was observed with deficit irrigation compared to full irrigation across both wheat and maize seasons. Compared to no manure under full irrigation, the application of manure under deficit irrigation showed an increase in leaf area index of 3.9% and 9.7% in wheat-Yr1 and wheat-Yr2, respectively. Meanwhile, the residual effect of manure with deficit irrigation resulted in an increase in leaf area index by 20.5% and 12.8% in maize-Yr1 and maize-Yr2, respectively. Manure application also resulted in higher root weight and root length density than treatments without manure (Figure 3), while full irrigation resulted in lower root weight and root length density than deficit irrigation. Compared to no manure under full irrigation, the application of manure under deficit irrigation resulted in a 14.4% increase in root length density and a 36.9% increase in root weight density in wheat Yr1. In maize Yr1, the residual effect of manure under deficit irrigation led to a 16.5% increase in root length density and a 2.7% increase in root weight density relative to no manure under full irrigation.

**Figure 3.**
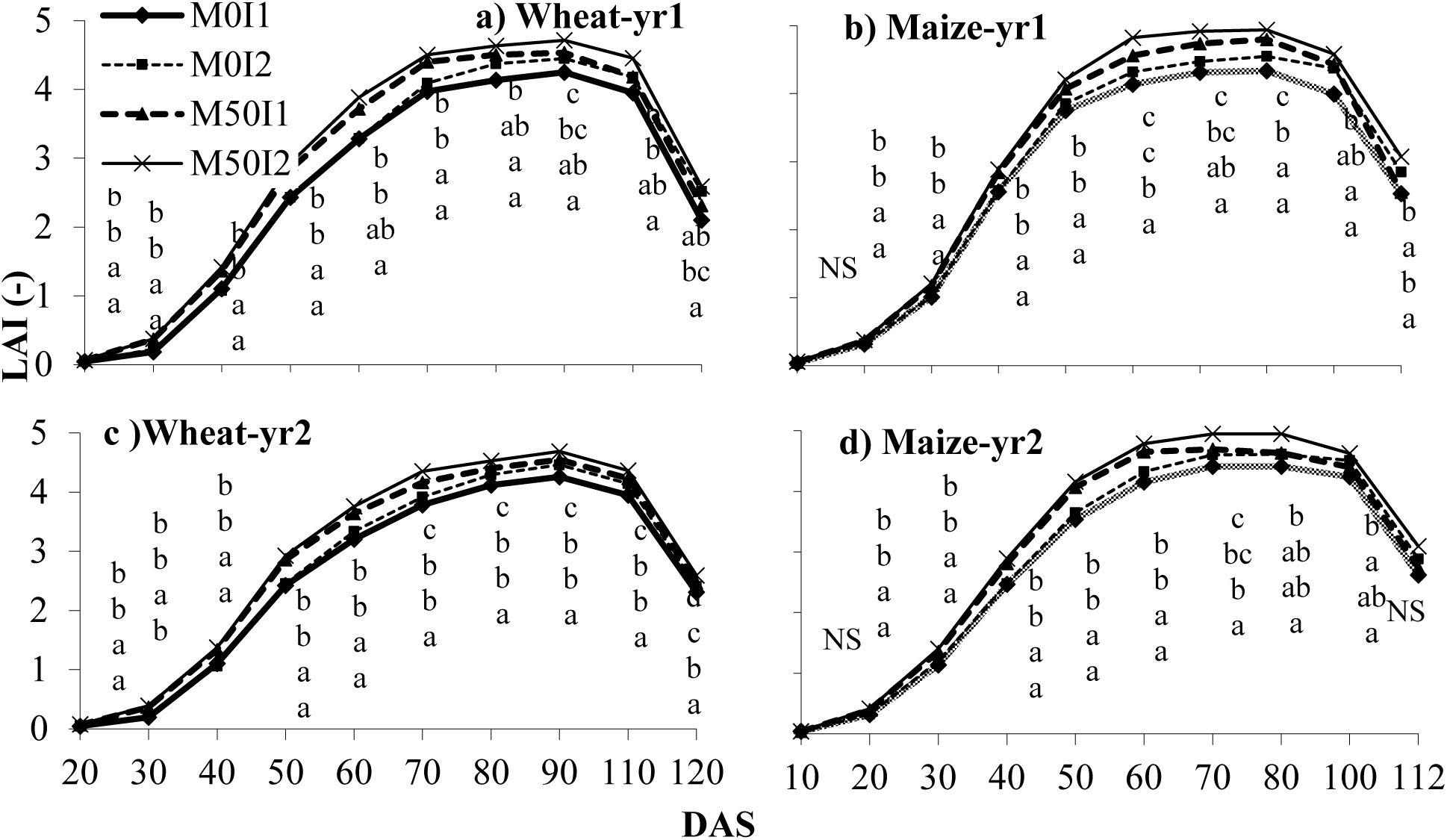
Effect of manure and irrigation on leaf area index measured at different time intervals during crop season.

**Figure 4.**
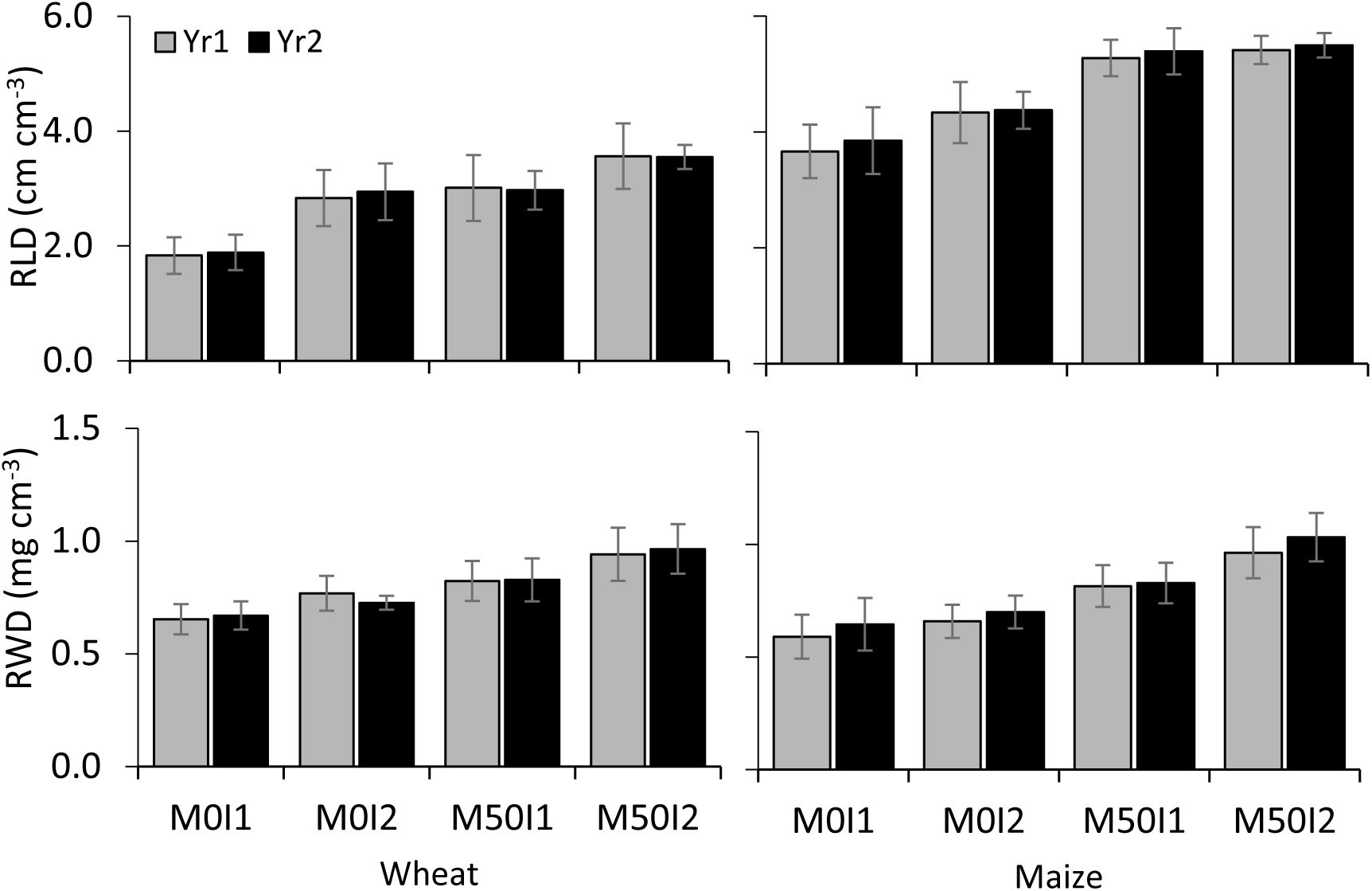
Effect of manure and irrigation levels on root length density and root weight density of wheat and maize

### 3.5. Bulk density, Infiltration rate, soil organic carbon, and Soil hydraulic conductivity

Soil bulk density increased with depth, with lower bulk density observed at shallower depths (0-5 cm) compared to deeper depths (20-30 cm). The application of manure resulted in a decrease in soil bulk density (Figure 5). However, the response to manure and irrigation was inconsistent across both maize and wheat seasons. In wheat (2008), at a depth of 0-5 cm, manure with full irrigation and manure with deficit irrigation decreased bulk density by 8.6% and 10.7%, respectively, compared to the 20-30 cm soil depth. In maize (2008), at a depth of 0-5 cm, manure with full irrigation and manure with deficit irrigation decreased bulk density by 10.1% and 7.6%, respectively, compared to the 20-30 cm soil depth. The application of manure resulted in an increase in the infiltration rate compared to scenarios without manure application (Figure 6; Figure S3). Greater infiltration rates were observed in both wheat seasons compared to both maize seasons. At 30 minutes of infiltration rate, in wheat Yr1, manure application with deficit irrigation increased the infiltration rate by 9.1% as compared to manure with full irrigation. The incorporation of manure alongside deficit irrigation led to an enhancement in hydraulic conductivity throughout both wheat and maize cultivation seasons (Figure 7). Hydraulic conductivity exhibited a higher value in the second year of both maize and wheat cultivation when compared to Yr1. Specifically, in the Yr1 of wheat cultivation, the combination of manure with deficit irrigation demonstrated an increase in hydraulic conductivity of 4.8%, whereas the residual effect of manure alongside deficit irrigation resulted in a 5.4% increase in maize cultivation’s Yr1. However, in the second year of wheat cultivation, deficit irrigation combined with manure decreased hydraulic conductivity by −8.5%. Conversely, there was no difference in the hydraulic conductivity values in Yr2 of maize cultivation compared to deficit irrigation paired with full irrigation. Soil organic carbon (SOC) was increased with application of manure as compared to no manure application (Figure 8). Deficit irrigation resulted in an increase in SOC compared to full irrigation. SOC decreased with increasing soil depth. In Yr2 of maize and wheat cultivation, SOC was higher as compared to Yr_1_ cultivation. At 0-5 cm soil depth, manure application with deficit irrigation increased SOC by 6.6% and 5.5% in wheat-Yr1 and wheat-Yr2, respectively. The residual effect of manure with deficit irrigation increased the soil organic matter by 5.7% and 1.5% in maize-Yr1 and maize-Yr2, respectively.

**Figure 5.**
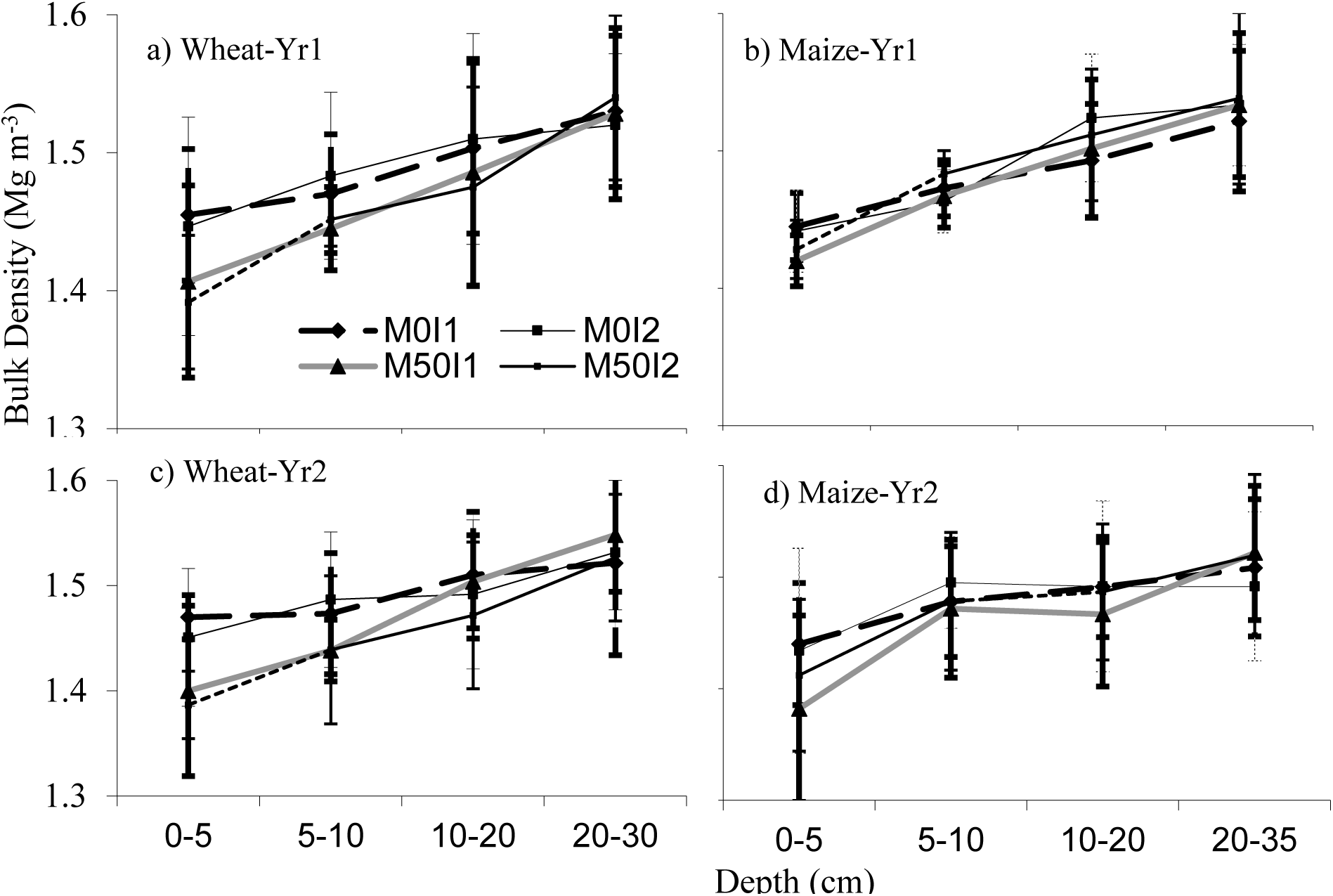
Effect of manure and irrigation levels on soil bulk density at different depths at harvest of wheat and maize

**Figure 6.**
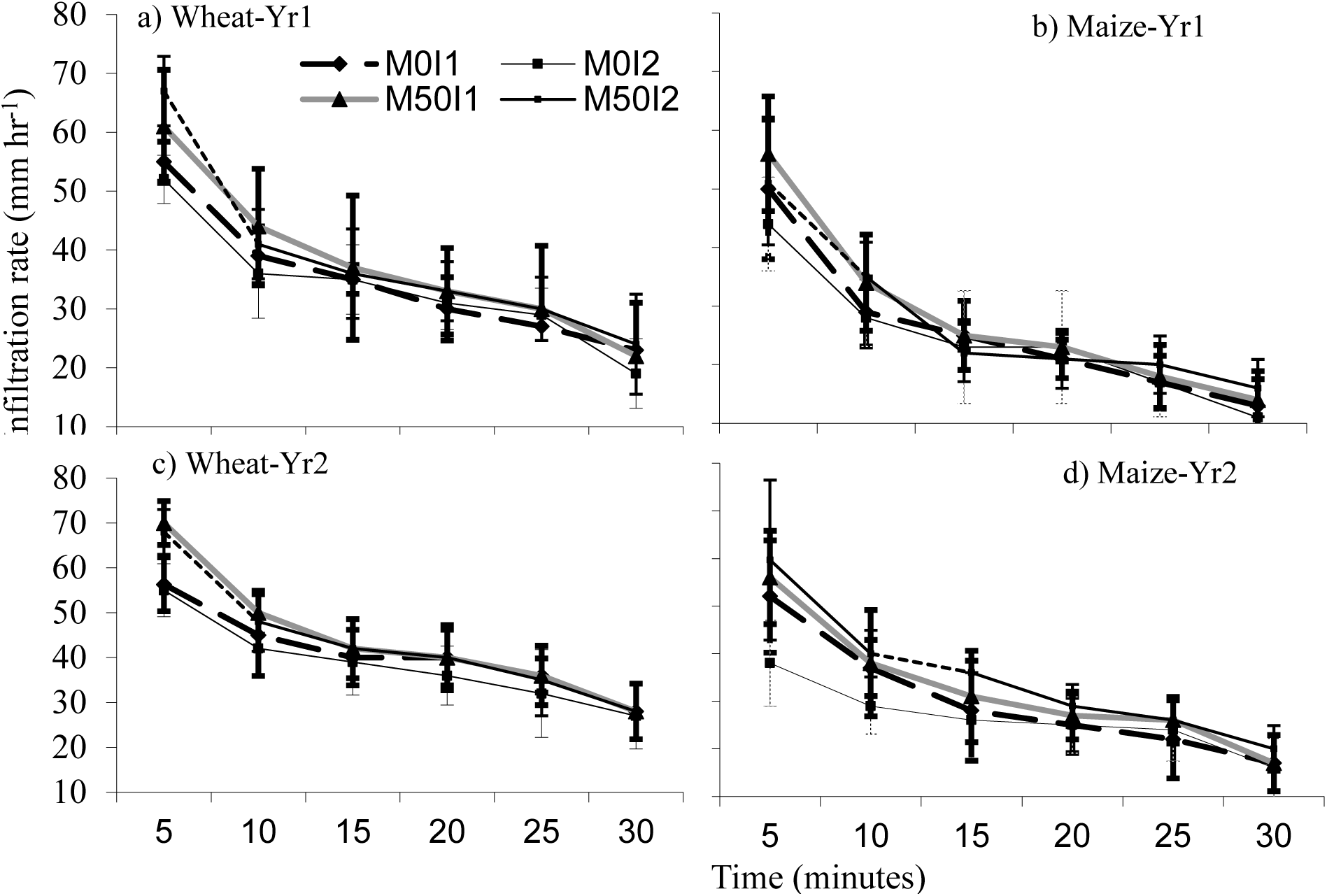
Effect of manure and irrigation levels on infiltration rate at harvest of a) wheat-1, b) maize-1, c) wheat-2, d) maize-2.

**Figure 7.**
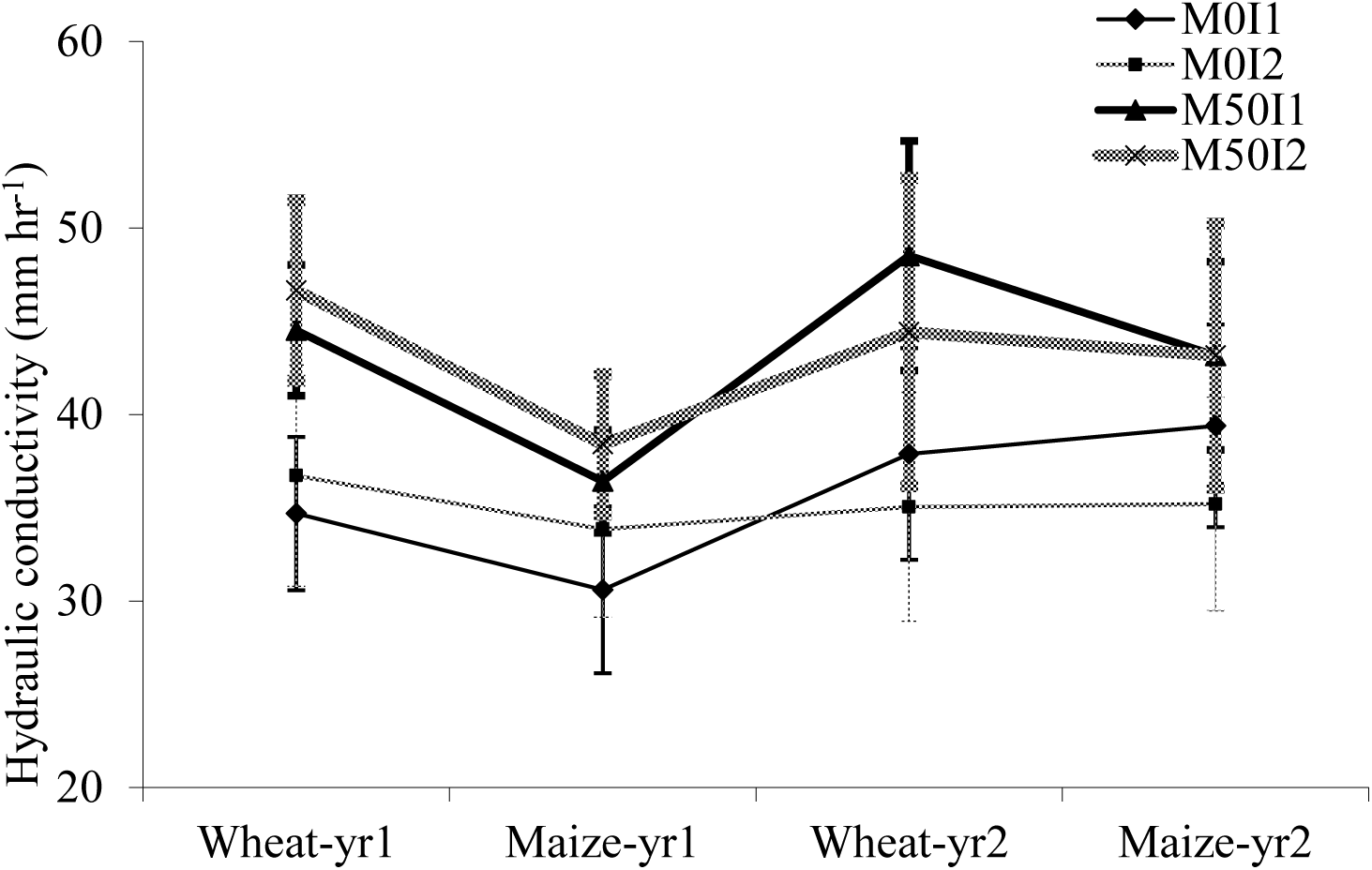
Soil saturated hydraulic conductivity under different treatments at the harvest of the crop

**Figure 8.**
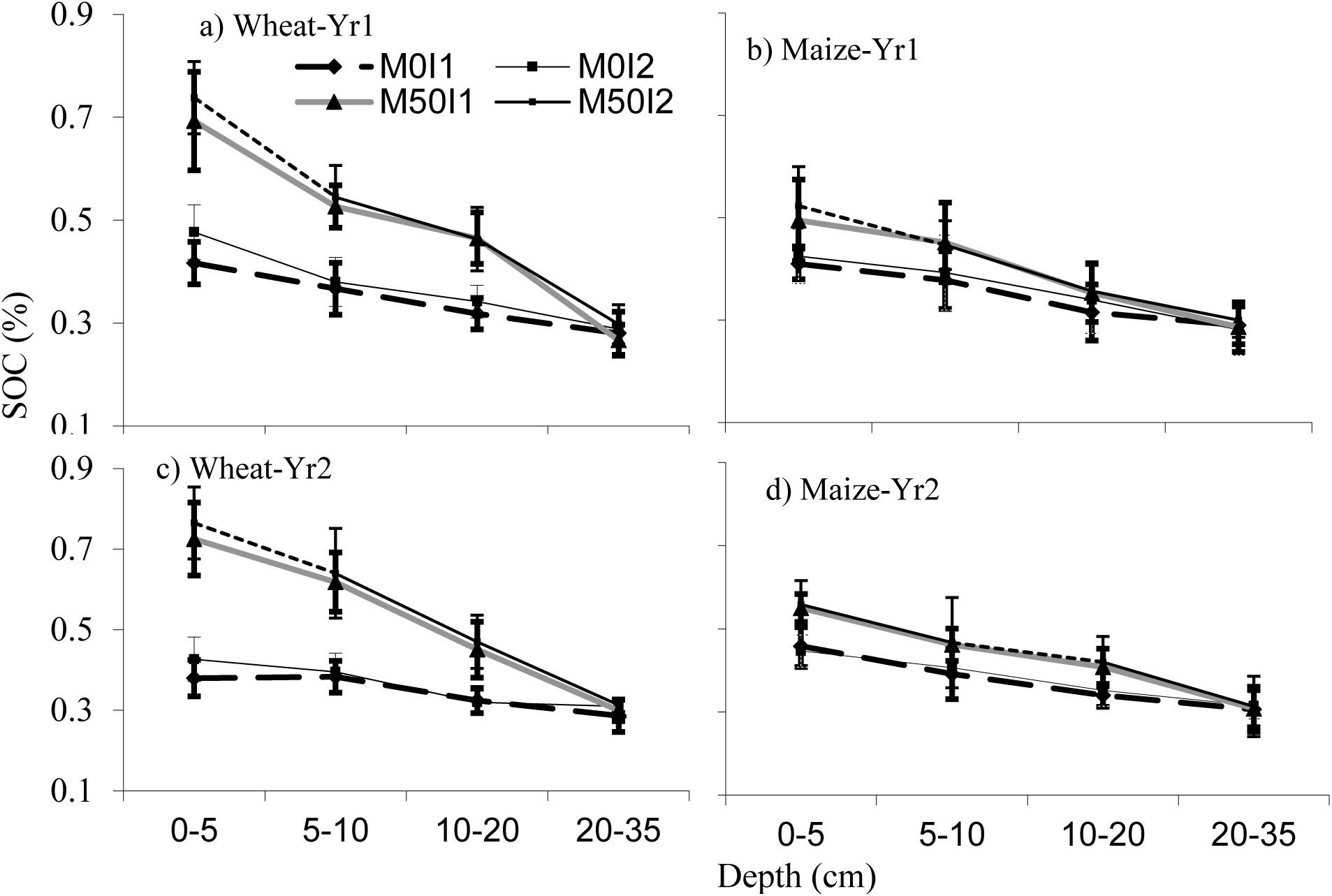
Effect of manure and irrigation levels on soil organic carbon at harvest of wheat and maize

### 3.6. Economic and Marginal Analysis

The economic analysis was performed for both of the growing seasons and the average values are presented in Table 8. The economic analysis conducted on annual basis (wheat + maize) proved that the combined application of manure and supplemental irrigation substantially improved the marginal economic returns of the wheat–maize system. Manure application resulted low VCR for wheat crop, however, its residual effect on succeeding maize crop increased the VCR, compared to non-manured treatments. The average annual VCR varied only from 1.34 to 1.42 among treatments, the marginal rate of return increased from 159.5% under M0×I2 to 205.1% under M50×I1 and reached 239.3% under M50×I2. Thus, the M50×I2 treatment provided the highest marginal economic return, indicating that investment in manure combined with the higher irrigation regime was economically attractive under the conditions of the study.

## 4. Discussion

### 4.1. Grain yield, irrigation water use efficiency, and water use efficiency

Manure application (M_50_) consistently enhanced grain yield and water use efficiency (WUE) for both wheat and maize compared to the control without manure (M_0_). These improvements align with previous findings that organic amendments enhance soil fertility, water retention, and nutrient availability, thereby improving overall crop productivity (Yagüe et al., 2016; (Yang et al., 2015; (Koutroubas et al., 2016). While full irrigation (I_2_) generally resulted in the highest yields, deficit irrigation (I_1_) combined with manure (M_50_I_1_) produced statistically comparable yields, particularly in maize. This suggests that the addition of organic matter can mitigate the adverse effects of moderate water stress, likely by improving root growth and the soil’s capacity to retain moisture. Evidently, the highest WUE and irrigation water use efficiency (WUE_i_) values were achieved under the M_50_I_1_ treatment, with maize WUE reaching 2.06 kg ha^1^ mm^1^ in the second season. These results indicate that manure improves water efficiency not just by enhancing soil structure and porosity, but also by facilitating better synchronization between nutrient release and plant demand. Research by (Eghball et al., 2004) has demonstrated that manure application can improve soil physical properties, leading to enhanced water infiltration and retention, which in turn can increase water use efficiency in crops. Under limited water availability, manure appears to serve as a compensatory mechanism, reducing non-productive water losses and maintaining crop yields. These findings highlight that the integration of manure with deficit irrigation can be a viable and sustainable strategy in semi-arid wheat-maize systems. The results are also consistent with previous studies (Yang et al., 2015; Tahir and Mulla, 2025), which reported that optimized nitrogen fertilization combined with suitable water management maintained high crop yields while significantly reducing nitrate leaching in wheat-maize rotations. Soil moisture data (Figure S2) indicated that deficit irrigation showed lower soil water contents at different intervals, compared to full irrigation, and below the critical limit of readily available water to plant, and thus showed soil water stress. Soil Water stress caused the decrease in grain yield of both crops.

### 4.2. Evapotranspiration, Fallow Evaporation, and Drainage Dynamics

The interaction between manure and irrigation significantly influenced crop evapotranspiration (ET_c_), though their main effects were non-significant. Full irrigation with manure (M_50_I_2_) led to the highest ET_c_ in both wheat and maize seasons, yet values under deficit irrigation with manure (M_50_I_1_) were statistically comparable, particularly in Yr_1_. This suggests that manure enhances water retention (Tahir et al., 2012) and compensates for reduced irrigation by improving soil physical properties, consistent with previous findings (Eghball et al., 2004; (Blanco-Canqui and Lal, 2007). Evaporation during the fallow period contributed 21.7% and 14.8% of annual ET_c_ in Yr1 and Yr2, respectively, highlighting a critical window of water loss. Drainage was highest under deficit irrigation without manure (M_0_I_1_), and consistently greater at 70 cm than at 110 cm soil depth. In contrast, manure application substantially reduced drainage under both irrigation regimes, aligning with findings that optimized management improves water and nitrogen use efficiency in wheat–maize systems (Singh et al., 2023; (Yang et al., 2015). In this study, manure applications significantly reduced drainage losses under both full and deficit irrigation, particularly at 70 cm depth. This effect is likely due to improved soil structure, increased water-holding capacity, and enhanced aggregation resulting from organic amendments (Blanco-Canqui and Lal, 2007). These improvements can limit percolation and promote more efficient use of irrigation water. Moreover, the interaction between manure and irrigation was statistically significant for ET_c_, with manure under full irrigation leading to the highest evapotranspiration. However, ET_c_ under deficit irrigation with manure remained comparable, indicating that manure can buffer the effects of limited water supply (Eghball et al., 2004).

### 4.3. Nitrate leaching and Nitrate build-up in soil

The interaction between manure application and irrigation significantly influenced nitrate (NO₃⁻ -N) leaching across both wheat and maize growing seasons, while their individual main effects were statistically nonsignificant. Manure application consistently increased nitrate leaching at both 70 cm and 110 cm soil depths, with more pronounced effects at 70 cm. For example, compared to the control under full irrigation, manure application under deficit irrigation led to a 65.9% and 91.4% increase in nitrate leaching in wheat (Yr1 and Yr2), and a 95.3% and 87.7% increase in maize (Yr1 and Yr2), respectively. These results are consistent with previous studies showing that the use of organic amendments like manure enhances nitrogen mineralization and elevates nitrate concentrations in the soil, particularly when water movement from irrigation or rainfall promotes leaching (Yang et al., 2015; Ju et al., 2006; Delin and Stenberg, 2014). Although deficit irrigation typically reduced total water percolation, nitrate concentrations in leachate remained elevated under manure treatments, suggesting asynchronous nutrient release and crop uptake. This interaction may be due to the mismatch between mineralization rates and crop nitrogen demand, increasing the risk of nitrate mobility during periods of reduced evapotranspiration and drainage accumulation (Ruidisch et al., 2013; Perego et al., 2012). Furthermore, although nitrate leaching at 70 cm depth was reduced in Yr2 compared to Yr1, it increased at 110 cm, suggesting downward movement of nitrate over time, likely driven by accumulated drainage and nitrogen accumulation in the soil profile. Nitrate buildup in the soil also showed a significant manure x irrigation interaction. Manure consistently led to higher NO₃^⁻^-N accumulation, especially 60 days after sowing, with levels decreasing by harvest. Full irrigation generally reduces nitrate buildup relative to deficit irrigation. At 60 days, manure under deficit irrigation led to increases of 8.0% and 19.3% in soil nitrate buildup in wheat-Yr1 and wheat-Yr2, respectively, compared to no manure with full irrigation. Residual effects in maize seasons showed smaller but remarkable increases (2.9-9.3%), suggesting carry-over of mineralized N between crops. These observations reflect the complex dynamics of nitrogen cycling in manured systems. While organic amendments enhance soil fertility and long-term productivity (Blanco-Canqui and Lal, 2007), they can also elevate nitrate leaching risks if not properly synchronized with plant uptake and irrigation scheduling. Efficient nitrogen management, including precise manure application timing and controlled irrigation, is essential to balance productivity and environmental sustainability (Gheysari et al., 2009; Cameron et al., 2013)

### 4.4. Root weight and root length density, Plant height, and leaf area index

The interaction between manure application and irrigation level had a significant influence on crop development indicators, including plant height, leaf area index (LAI), and root system traits, although neither factor alone had a strong effect. Across both wheat and maize growing seasons, manure application under deficit irrigation (M_50_I_1_) led to significantly greater plant height at all growth stages compared to the other treatment combinations (Fig. 2). At 106 DAS, plant height increased slightly (1.3% in maize-Yr1 and 0.9% in maize-Yr2) with manure under deficit irrigation, highlighting manure’s role in supporting plant growth under limited water through improved nutrient availability and soil biology (Arif et al., 2016; Blanco-Canqui and Lal, 2007). The LAI also showed a significant manure and irrigation interaction effect (Fig. 3). Highest LAI values were consistently recorded under M_50_I_1_ across all growth stages in both crops. Compared to M_0_I_2_, manure under deficit irrigation increased LAI by 3.9% and 9.7% in wheat-Yr1 and wheat-Yr2, respectively, and by 20.5% and 12.8% in maize-Yr1 and maize-Yr2, respectively. These results support the idea that organic amendments promote canopy growth by boosting root absorption and better coordination of water and nutrient availability (Eghball et al., 2004; (Bhatt et al., 2016). Root development followed a similar trend, with M_50_I_1_ showing significantly higher root weight and root length density compared to all other treatments (see Fig. 4). In wheat-Yr1, manure with deficit irrigation resulted in a 36.9% increase in root weight and a 14.4% increase in root length density relative to M_0_I_2_. The residual effect of manure in maize-Yr1 also enhanced root traits, with 16.5% greater root length density and 2.7% higher root weight density under M_50_I_1_ compared to M_0_I_2_. These results highlight the collaborative effect of organic inputs and moderate water stress in promoting deeper and denser root systems, key behaviors for water acquisition in semi-arid cropping systems (Fageria, 2012; Bello et al., 2021). Climatic conditions during the study period (Fig. 1) revealed considerable variability in precipitation, ET₀, and temperature patterns, which likely influenced soil moisture dynamics and root responses. Higher wind speeds and evapotranspiration during key growth phases would have intensified crop water demand, particularly under deficit irrigation. In such scenarios, the role of organic amendments becomes even more critical in enhancing root-soil interactions and buffering against climatic stress. Overall, the integrated application of manure with deficit irrigation improved crop physiological parameters and resource use efficiency by stimulating both above- and below-ground biomass. These improvements are essential for sustainable intensification in water-limited environments.

### 4.5. Bulk density, Infiltration rate, soil organic carbon, and Soil hydraulic conductivity

The combined effects of manure application and irrigation significantly influenced key soil physical properties like bulk density, infiltration rate, hydraulic conductivity, and soil organic carbon (SOC). However, responses varied across seasons and depths. Soil bulk density decreased at surface layers (0-5 cm) compared to deeper layers, particularly under manure treatments. In wheat-Yr_1_, manure combined with deficit irrigation (M_50_I_1_) reduced bulk density by 10.7% at the surface, supporting findings that organic inputs enhance soil aggregation and reduce compaction in the root zone (Blanco-Canqui and Lal, 2007; Rengel, 2011). Infiltration rates were consistently higher in manure-amended plots (Fig. 6). While full irrigation generally promoted greater infiltration, the combination of manure with deficit irrigation (M_50_I_1_) yielded the highest infiltration in wheat-Yr1 and maize-Yr1, with an increase of 9.1% over full irrigation. This suggests that organic matter improves water entry into the soil, especially under reduced irrigation, by enhancing macropore continuity and surface stability (Bello et al., 2021; Ghimire et al., 2017). Soil hydraulic conductivity showed a similar trend. Manure with deficit irrigation increased conductivity by 4.8% in wheat-Yr1 and 5.4% in maize-Yr1, indicating improved soil pore structure and water transmission. However, this positive effect was not observed in Yr2 wheat, where conductivity declined slightly. This study indicated that the application of organic amendments enhanced plant water use efficiency and improved soil structure, leading to better maize growth and yield (Sisouvanh et al., 2021). Soil organic carbon (SOC) increased notably with manure application, particularly in the upper 0-5 cm soil layer. The greatest SOC values were recorded under manure combined with deficit irrigation, with increases of 6.6% in wheat-Yr1 and 5.7% in maize-Yr1 compared to the no manure control (Fig. 8). Deficit irrigation may have contributed to higher SOC by slowing decomposition rates and promoting microbial carbon stabilization, consistent with previous findings (Chivenge et al., 2011). Additionally, SOC levels decreased with soil depth, likely due to the surface application of manure and limited downward movement of organic materials. These findings highlight the value of combining manure application with deficit irrigation to enhance soil structure, increase water-holding capacity, and boost carbon sequestration, key factors for promoting sustainability and resilience in semi-arid agricultural systems.

### 4.6. Economics of irrigation and manure applications

Economic analysis (Table 7) is ultimate yardstick to judge a treatment effectively in terms of economic analysis and to recommend a specific technology. Applying manure was not economically effective when considered for only one crop as it decreased the VCR. However, proceeding maize crop showed higher VCR under the residual effect of manure. Overall, manure application with irrigation at 100% ET_c_ showed highest marginal rate of return (MRR). Deficit irrigation at 75% ET_c_ with manure was second best option. Deficit irrigation without manure was totally not a feasible option. Tahir et al. (2026) developed a relationship between irrigation and yield, and stated that better irrigation scheduling enhances the yield, but excessive irrigation could not be cost effective as it decreases the crop yield. Higher MRR observed with manure under full irrigation was a result of better availability of nutrients with optimum soil moisture under full irrigation. Si et al. (2021)] found that optimum nitrogen application yields highest mean return as compared to other nitrogen levels.

**Table 7.**
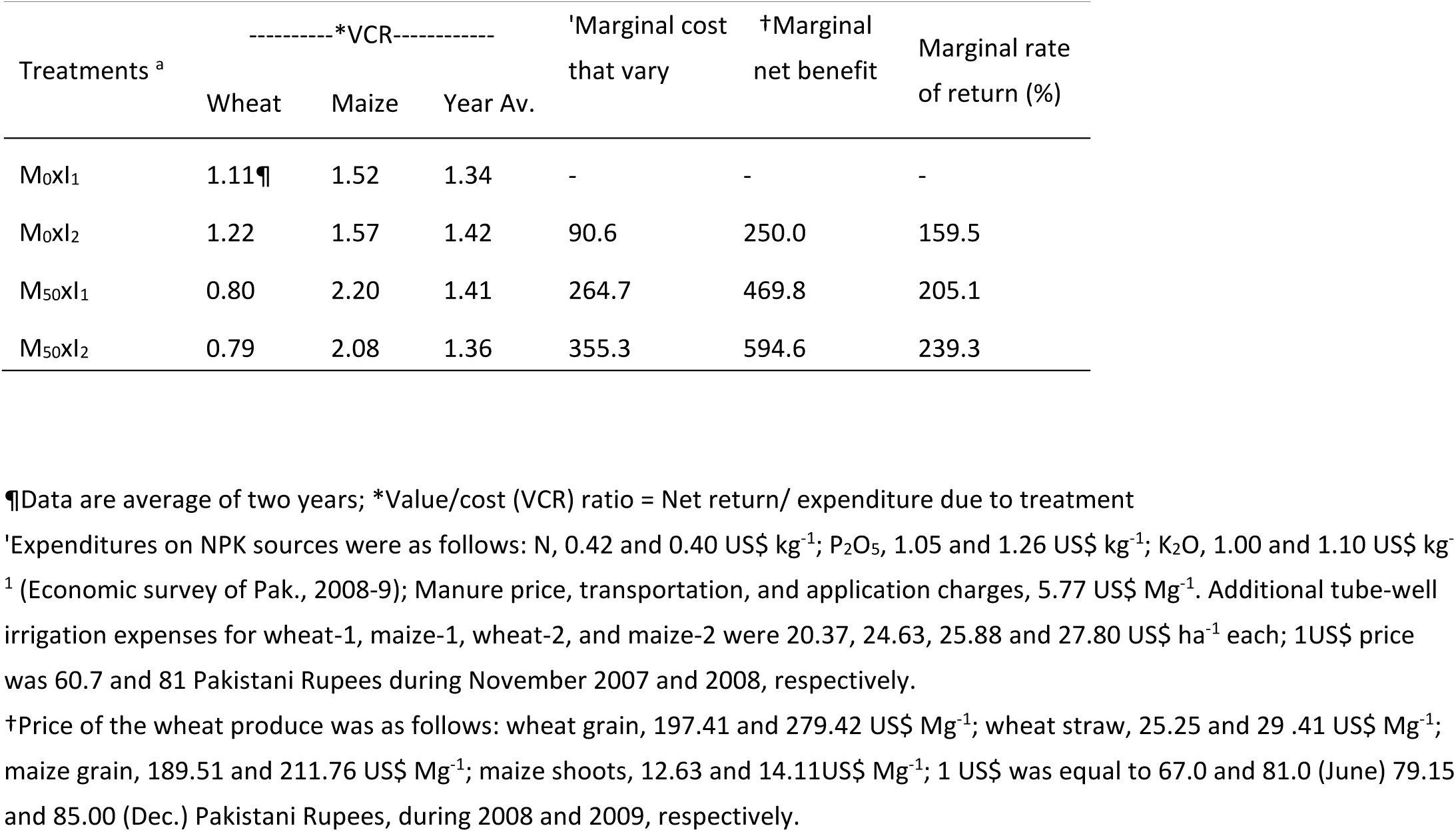
Economic analysis of wheat and maize under different treatment combinations of manure and irrigation.

| Treatments <sup>a</sup> | -----*VCR----- |  |  | 'Marginal cost<br>that vary | †Marginal<br>net benefit | Marginal rate<br>of return (%) |
| --- | --- | --- | --- | --- | --- | --- |
|  | Wheat | Maize | Year Av. |  |  |  |
| M<sub>0</sub>XI<sub>1</sub> | 1.11¶ | 1.52 | 1.34 | - | - | - |
| M<sub>0</sub>XI<sub>2</sub> | 1.22 | 1.57 | 1.42 | 90.6 | 250.0 | 159.5 |
| M<sub>50</sub>XI<sub>1</sub> | 0.80 | 2.20 | 1.41 | 264.7 | 469.8 | 205.1 |
| M<sub>50</sub>XI<sub>2</sub> | 0.79 | 2.08 | 1.36 | 355.3 | 594.6 | 239.3 |

¶Data are average of two years; \*Value/cost (VCR) ratio = Net return/ expenditure due to treatment
<sup>†</sup>Expenditures on NPK sources were as follows: N, 0.42 and 0.40 US\$ kg<sup>-1</sup>; P<sub>2</sub>O<sub>5</sub>, 1.05 and 1.26 US\$ kg<sup>-1</sup>; K<sub>2</sub>O, 1.00 and 1.10 US\$ kg<sup>-1</sup> (Economic survey of Pak., 2008-9); Manure price, transportation, and application charges, 5.77 US\$ Mg<sup>-1</sup>. Additional tube-well irrigation expenses for wheat-1, maize-1, wheat-2, and maize-2 were 20.37, 24.63, 25.88 and 27.80 US\$ ha<sup>-1</sup> each; 1US\$ price was 60.7 and 81 Pakistani Rupees during November 2007 and 2008, respectively.
†Price of the wheat produce was as follows: wheat grain, 197.41 and 279.42 US\$ Mg<sup>-1</sup>; wheat straw, 25.25 and 29.41 US\$ Mg<sup>-1</sup>; maize grain, 189.51 and 211.76 US\$ Mg<sup>-1</sup>; maize shoots, 12.63 and 14.11US\$ Mg<sup>-1</sup>; 1 US\$ was equal to 67.0 and 81.0 (June) 79.15 and 85.00 (Dec.) Pakistani Rupees, during 2008 and 2009, respectively.

## 5. Conclusions

The findings of this study underscore the importance of integrating organic amendments with irrigation management to enhance the productivity and sustainability of wheat–fallow–maize systems in semi-arid regions. Combining farmyard manure with deficit irrigation emerged as an effective approach, achieving yields like full irrigation while enhancing water use efficiency and lowering total water requirements. Manure application positively influenced plant height, leaf area index, and root development, contributing to better crop performance under reduced water input. It also improved soil physical properties, including reduced bulk density, increased infiltration, and enhanced hydraulic conductivity. The increase in soil organic carbon, especially in the topsoil, signals potential long-term enhancements in soil quality. However, the benefits of manure were accompanied by increased nitrate leaching, especially during the fallow period and deeper soil layers over time. This highlights the need for more effective nutrient management approaches, such as improving the timing of manure application or incorporating cover crops, to reduce potential environmental impacts. In conclusion, combining manure with deficit irrigation is an effective approach to enhance yield, conserve water, and improve soil health. However, careful management is needed to minimize nitrate leaching. These findings provide practical insights for sustainable farming in water-limited regions and highlight the need for further research on reducing nutrient losses.

## Supporting information

Yield Response and Nitrate Leaching_Sfile

## Declarations

### Availability of data and materials

Data will be available upon request. The original contributions presented in this study are included in the article/supplementary material. Further inquiries can be directed to the corresponding author(s).

## Acknowledgments

The authors appreciate Dr. Abdul Ghaffar Niazi’s help in collecting some field data.

## Supplementary Materials

The following supporting information is provided with manuscript, Table S1. Physical and chemical characteristics of the soil of the experimental site; Table S2.Properties of dairy manure used for field Trial; Table S3. Soil sampling and leachate collection schedule for NO_3_-N; Table S4. Water use efficiency of wheat and maize crops during different years under different irrigation and manure treatments; Table S5. Treatment effect on Root Length Density; Figure S1. Wheat trial and leachates collection, and measurement of soil hydraulic properties under field conditions; Figure S2. Moisture measured before each irrigation, and critical limit of readily available water; Figure S3. Effect of manure and irrigation levels on soil saturated hydraulic conductivity at harvest of wheat and maize.

## Authors’ contributions

Conceptualization: Muhammad Tahir, Anwar ul Hasssan, and David Mulla; methodology, software, validation, and formal analysis: Muhammad Tahir, Daavid Mulla, and Saliha Maqbool; investigation, resources, data curation: Saliha Maqbool and Muhammad Tahir; Writing—original draft preparation: Muhammad Tahir, Saliha Maqbool; Writing—review and editing; Mu-hamad Tahir, David Mulla; visualization, supervision, and project administration: Muhammad Tahir and Anwar Ul Hassan; funding acquisition, Muhammad Tahir, and Anwar Ul Hassan. All authors have read and agreed to the published version of the manuscript.

## Abbreviations

The following abbreviations are used in this manuscript:

WUE_i_: irrigation water use efficiency
N: Nitrogen
SOM: soil organic matter
SOC: soil organic carbon
P: phosphorus
K: potash
GY: grain yield
LAI: leaf area index
K_fs_: Soil saturated hydraulic conductivity
DAS: days after sowing
RWD: Root weight density
RLD: Root length density
WUE: water use efficiency
ET_c_: crop evapotranspiration
LSD: least significant difference
VCR: Value cost ratio

