## Supplementary material for "Field Modeling Study of Yield Response and Nitrate Leaching with Manure Application and Deficit Irrigation for Maize-Fallow-Wheat Rotation": Yield Response and Nitrate Leaching_Sfile

Appendix

**Table S1 Physical and chemical characteristics of the soil of the experimental site**

| **Characteristics** | | **Unit** | **Value** |
| --- | --- | --- | --- |
| Sand: Silt: Clay | | % | 38:37:25 |
| Textural Class (upper 15cm) | | Loam | |
| Bulk density (0-5, 5-10, and 10-20 cm |  | Mg m^-3^ | 1.43, 1.45, 1.49 |
| Infiltration rate | | mm hr^-1^ | 46.5 |
| Field saturated hydraulic conductivity  (upper 15cm) | | " | 54.4 |
| Penetration resistance at 15 % water content  (upper 15cm) | | kPa | 1025 |
| EC_e_ | | dS m^-1^ | 1.67 |
| pH | |  | 8.0 |
| Soil organic carbon | 0-5 cm | g kg^-1^ | 5.2 |
|  | 5-10 cm |  | 4.4 |
|  | 10-20 cm |  | 3.6 |
| Total nitrogen | 0-15 cm | g kg^-1^ | 0.47 |
|  | 15-30 cm |  | 0.35 |
| Available phosphorus | 0-15 cm | mg kg^-1^ | 8.4 |
|  | 15-30 cm |  | 7.2 |
| Available potassium | 0-15 cm | " | 115.4 |
|  | 15-30 cm |  | 97.2 |
| NO_3_^-^ | 0-10 cm | " | 6.62 |
|  | 10-20 cm | " | 7.43 |
|  | 20-40 cm | " | 6.82 |
|  | 40-60 cm | " | 6.12 |
|  | 60-80 cm | " | 5.64 |
|  | 80-115 cm | " | 5.03 |

**Table S2.** Properties of dairy manure used for field Trial.

| Parameter | Unit | Year-1 | Year-2 |
| --- | --- | --- | --- |
| Moisture contents | (%) | 60.5 ± 2.50* | 68.4 ± 3.30 |
| N | - | 1.42 ± 0.02 | 1.38 ± 0.03 |
| P_2_O_5_ | - | 0.47 ± 0.02 | 0.50 ± 0.01 |
| K_2_O | - | 1.30 ± 0.04 | 1.20 ± 0.03 |

*, manure amended soil; ¶, means± standard deviation

| **^Sampling No. with crop^** | **Soil Sampling** | | | **Leachate collection** | |
| --- | --- | --- | --- | --- | --- |
|  | DAS | No. of Irrigations up to sampling | | No of Irrigations when leachate collected | |
|  |  | I_1_ | I_2_ | I_1_ | I_2_ |
| 1 (Wheat) | 60 | 2 | 2 | 1 | 1 |
| 2 - | 90 | 3 | 4 | 2 | 2 |
| 3 - | harvest | 4 | 6 | 3 | 4 |
| 4 - | - | - | - | 4 | 6 |
| Fallow | End period (last week of fallow period) | | | | |
| 1 (Maize) | 30 | 2 | 2 | 1 | 1 |
| 2 - | 60 | 4 | 5 | 2 | 2 |
| 3 - | harvest | 6 | 8 | 4 | 5 |
| 4 - | - | - | - | 6 | 8 |

**Table S3:** Soil sampling and leachates collection schedule for NO_3_-N

**Table S4:** Water use efficiency of wheat and maize crop during different years under different irrigation and manure treatments.

|  | WUE (kg ha^-1^ mm^-1^) | | | |
| --- | --- | --- | --- | --- |
|  | Wheat-1 | Wheat-2 | Maize-1 | Maize-2 |
| M_0_* | 0.80 | 0.74 | 1.44 | 1.66 |
| M_50_ | 0.93 | 0.97 | 1.84 | 1.97 |
| I_1_** | 0.90 | 0.92 | 1.61 | 1.50 |
| I_2_ | 0.92 | 0.92 | 1.50 | 1.49 |
| M_0_I_1_ | 0.82 b | 0.70 c | 1.27 b | 1.44 b |
| M_0_I_2_ | 0.78 b | 0.77 b | 1.61 a | 1.87 a |
| M_50_I_1_ | 0.93 a | 0.98 a | 1.82 a | 1.88 a |
| M_50_I_2_ | 0.93 a | 0.96 a | 1.85 a | 2.06 a |
| LSD (p≤ 0.05) M | 0.07 | 0.1 | 0.2 | 0.19 |
| I | NS | N. S | NS | 0.22 |
| M x I | 0.06,0.09 | 0.05,0.11 | 0.30,0.29 | 0.31,0.29 |

**Table s5.** Treatment effect on Root Length Density

| **LSD (**p≤**0.05)** | **Wheat-1** | **Wheat-2** | **Maize-1** | **Maize-2** |
| --- | --- | --- | --- | --- |
| **RWD** | | | | |
| M | 0.12 | 0.12 | 0.32 | 0.11 |
| I | 0.08 | 0.05 | N.S. | 0.20 |
| M x I* | 0.11, 0.14 | 0.08, 0.13 | 0.17, 0.15 | 0.15, 0.16 |
| **RLD** | | | | |
| M | 0.17 | 0.22 | 0.51 | 0.46 |
| I | 0.29 | 0.31 | 0.21 | 0.49 |
| M x I | 0.42, 0.34 | 0.25, 0.52 | 0.31, 0.56 | 0.58, 52 |

RWD, root weight density; RLD, root length density; * 1^st^ LSD value is for same levels of Manure, while 2^nd^ for different levels of manure; Note: year effect was statistically non-significant for both crops

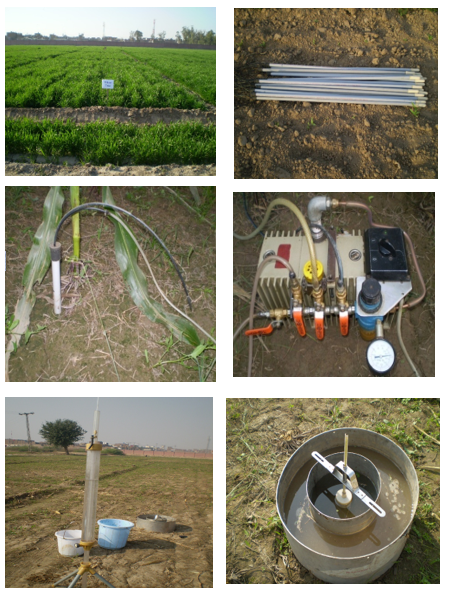

**Figure S1** Wheat trial and leachates collection, and measurement of soil hydraulic properties under field conditions

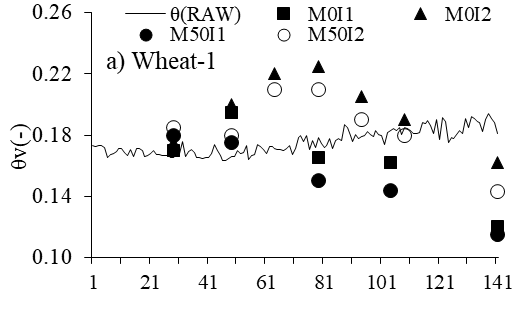

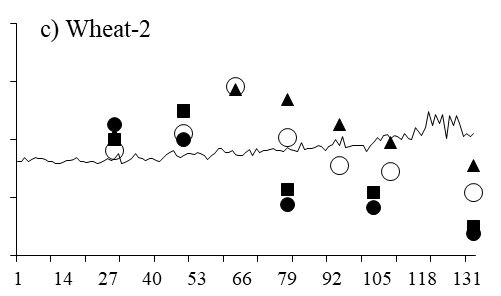

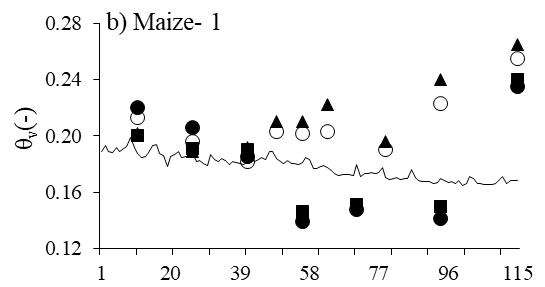

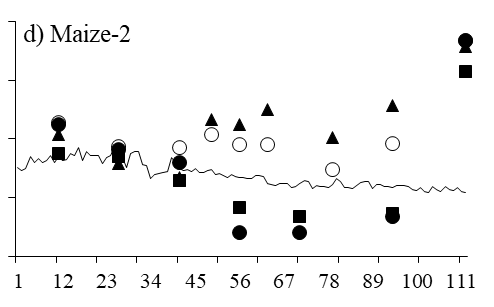

a) Wheat-1

θv(-)

|  | θ(RAW) | M0I1 | M0I2 | M50I1 | M50I2 |
| --- | --- | --- | --- | --- | --- |
| 1 | 0.173194 |  |  |  |  |
| 2 | 0.172354 |  |  |  |  |
| 3 | 0.173305 |  |  |  |  |
| 4 | 0.173113 |  |  |  |  |
| 5 | 0.172 |  |  |  |  |
| 6 | 0.165235 |  |  |  |  |
| 7 | 0.167332 |  |  |  |  |
| 8 | 0.167993 |  |  |  |  |
| 9 | 0.169441 |  |  |  |  |
| 10 | 0.170971 |  |  |  |  |
| 11 | 0.171259 |  |  |  |  |
| 12 | 0.168377 |  |  |  |  |
| 13 | 0.166604 |  |  |  |  |
| 14 | 0.170306 |  |  |  |  |
| 15 | 0.165666 |  |  |  |  |
| 16 | 0.171202 |  |  |  |  |
| 17 | 0.171142 |  |  |  |  |
| 18 | 0.17003 |  |  |  |  |
| 19 | 0.165936 |  |  |  |  |
| 20 | 0.166316 |  |  |  |  |
| 21 | 0.167716 |  |  |  |  |
| 22 | 0.169929 |  |  |  |  |
| 23 | 0.167448 |  |  |  |  |
| 24 | 0.167358 |  |  |  |  |
| 25 | 0.166865 |  |  |  |  |
| 26 | 0.166684 |  |  |  |  |
| 27 | 0.166805 |  |  |  |  |
| 28 | 0.165203 |  |  |  |  |
| 29 | 0.165508 | 0.17 | 0.175 | 0.18 | 0.185 |
| 30 | 0.16731 |  |  |  |  |
| 31 | 0.167936 |  |  |  |  |
| 32 | 0.175731 |  |  |  |  |
| 33 | 0.165634 |  |  |  |  |
| 34 | 0.169052 |  |  |  |  |
| 35 | 0.170771 |  |  |  |  |
| 36 | 0.168479 |  |  |  |  |
| 37 | 0.165137 |  |  |  |  |
| 38 | 0.165076 |  |  |  |  |
| 39 | 0.164662 |  |  |  |  |
| 40 | 0.165213 |  |  |  |  |
| 41 | 0.165061 |  |  |  |  |
| 42 | 0.167898 |  |  |  |  |
| 43 | 0.173525 |  |  |  |  |
| 44 | 0.169818 |  |  |  |  |
| 45 | 0.166786 |  |  |  |  |
| 46 | 0.163244 |  |  |  |  |
| 47 | 0.163572 |  |  |  |  |
| 48 | 0.164698 |  |  |  |  |
| 49 | 0.165921 | 0.195 | 0.2 | 0.175 | 0.18 |
| 50 | 0.165985 |  |  |  |  |
| 51 | 0.170159 |  |  |  |  |
| 52 | 0.168036 |  |  |  |  |
| 53 | 0.168882 |  |  |  |  |
| 54 | 0.173051 |  |  |  |  |
| 55 | 0.164245 |  |  |  |  |
| 56 | 0.166722 |  |  |  |  |
| 57 | 0.167022 |  |  |  |  |
| 58 | 0.17356 |  |  |  |  |
| 59 | 0.172227 |  |  |  |  |
| 60 | 0.170556 |  |  |  |  |
| 61 | 0.167798 |  |  |  |  |
| 62 | 0.172558 |  |  |  |  |
| 63 | 0.172635 |  |  |  |  |
| 64 | 0.17103 |  | 0.22 |  | 0.21 |
| 65 | 0.170489 |  |  |  |  |
| 66 | 0.170562 |  |  |  |  |
| 67 | 0.169983 |  |  |  |  |
| 68 | 0.170873 |  |  |  |  |
| 69 | 0.173032 |  |  |  |  |
| 70 | 0.167421 |  |  |  |  |
| 71 | 0.170914 |  |  |  |  |
| 72 | 0.178354 |  |  |  |  |
| 73 | 0.179486 |  |  |  |  |
| 74 | 0.175882 |  |  |  |  |
| 75 | 0.179445 |  |  |  |  |
| 76 | 0.170827 |  |  |  |  |
| 77 | 0.176433 |  |  |  |  |
| 78 | 0.172095 |  |  |  |  |
| 79 | 0.178698 | 0.165 | 0.225 | 0.15 | 0.21 |
| 80 | 0.174313 |  |  |  |  |
| 81 | 0.171935 |  |  |  |  |
| 82 | 0.172834 |  |  |  |  |
| 83 | 0.177611 |  |  |  |  |
| 84 | 0.171089 |  |  |  |  |
| 85 | 0.176301 |  |  |  |  |
| 86 | 0.177608 |  |  |  |  |
| 87 | 0.176376 |  |  |  |  |
| 88 | 0.187006 |  |  |  |  |
| 89 | 0.185113 |  |  |  |  |
| 90 | 0.179752 |  |  |  |  |
| 91 | 0.175545 |  |  |  |  |
| 92 | 0.180621 |  |  |  |  |
| 93 | 0.177878 |  |  |  |  |
| 94 | 0.179005 |  | 0.205 |  | 0.19 |
| 95 | 0.179792 |  |  |  |  |
| 96 | 0.182287 |  |  |  |  |
| 97 | 0.180096 |  |  |  |  |
| 98 | 0.17889 |  |  |  |  |
| 99 | 0.182653 |  |  |  |  |
| 100 | 0.180043 |  |  |  |  |
| 101 | 0.179603 |  |  |  |  |
| 102 | 0.173666 |  |  |  |  |
| 103 | 0.181586 |  |  |  |  |
| 104 | 0.183175 | 0.162 |  | 0.144 |  |
| 105 | 0.184524 |  |  |  |  |
| 106 | 0.182859 |  |  |  |  |
| 107 | 0.184894 |  |  |  |  |
| 108 | 0.180281 |  |  |  |  |
| 109 | 0.185179 |  | 0.19 |  | 0.18 |
| 110 | 0.184416 |  |  |  |  |
| 111 | 0.182663 |  |  |  |  |
| 112 | 0.181078 |  |  |  |  |
| 113 | 0.18077 |  |  |  |  |
| 114 | 0.181923 |  |  |  |  |
| 115 | 0.188047 |  |  |  |  |
| 116 | 0.187664 |  |  |  |  |
| 117 | 0.189561 |  |  |  |  |
| 118 | 0.185924 |  |  |  |  |
| 119 | 0.180863 |  |  |  |  |
| 120 | 0.190041 |  |  |  |  |
| 121 | 0.177045 |  |  |  |  |
| 122 | 0.191651 |  |  |  |  |
| 123 | 0.188794 |  |  |  |  |
| 124 | 0.175231 |  |  |  |  |
| 125 | 0.178505 |  |  |  |  |
| 126 | 0.177601 |  |  |  |  |
| 127 | 0.18002 |  |  |  |  |
| 128 | 0.182785 |  |  |  |  |
| 129 | 0.180824 |  |  |  |  |
| 130 | 0.190501 |  |  |  |  |
| 131 | 0.18497 |  |  |  |  |
| 132 | 0.191867 |  |  |  |  |
| 133 | 0.190737 |  |  |  |  |
| 134 | 0.188834 |  |  |  |  |
| 135 | 0.188448 |  |  |  |  |
| 136 | 0.182098 |  |  |  |  |
| 137 | 0.18993 |  |  |  |  |
| 138 | 0.193898 |  |  |  |  |
| 139 | 0.190649 |  |  |  |  |
| 140 | 0.187997 |  |  |  |  |
| 141 | 0.18095 | 0.12 | 0.162 | 0.115 | 0.143 |

c) Wheat-2

|  | θ(RAW) | M0I1 | M0I2 | M50I1 | M50I2 |
| --- | --- | --- | --- | --- | --- |
| 1 | 0.164553 |  |  |  |  |
| 2 | 0.164337 |  |  |  |  |
| 3 | 0.167389 |  |  |  |  |
| 4 | 0.164519 |  |  |  |  |
| 5 | 0.165949 |  |  |  |  |
| 6 | 0.167075 |  |  |  |  |
| 7 | 0.166692 |  |  |  |  |
| 8 | 0.166358 |  |  |  |  |
| 9 | 0.166244 |  |  |  |  |
| 10 | 0.16468 |  |  |  |  |
| 11 | 0.164781 |  |  |  |  |
| 12 | 0.162964 |  |  |  |  |
| 13 | 0.1634 |  |  |  |  |
| 14 | 0.163897 |  |  |  |  |
| 15 | 0.16513 |  |  |  |  |
| 16 | 0.165541 |  |  |  |  |
| 17 | 0.165847 |  |  |  |  |
| 18 | 0.167431 |  |  |  |  |
| 19 | 0.164621 |  |  |  |  |
| 20 | 0.164781 |  |  |  |  |
| 21 | 0.164026 |  |  |  |  |
| 22 | 0.16468 |  |  |  |  |
| 23 | 0.164509 |  |  |  |  |
| 24 | 0.163099 |  |  |  |  |
| 25 | 0.164011 |  |  |  |  |
| 26 | 0.165385 |  |  |  |  |
| 27 | 0.166799 |  |  |  |  |
| 28 | 0.165539 |  |  |  |  |
| 29 | 0.166798 | 0.18 | 0.183 | 0.19 | 0.172 |
| 30 | 0.169844 |  |  |  |  |
| 31 | 0.163025 |  |  |  |  |
| 32 | 0.164232 |  |  |  |  |
| 33 | 0.16535 |  |  |  |  |
| 34 | 0.166611 |  |  |  |  |
| 35 | 0.170251 |  |  |  |  |
| 36 | 0.16769 |  |  |  |  |
| 37 | 0.166824 |  |  |  |  |
| 38 | 0.165484 |  |  |  |  |
| 39 | 0.166766 |  |  |  |  |
| 40 | 0.16676 |  |  |  |  |
| 41 | 0.165014 |  |  |  |  |
| 42 | 0.16474 |  |  |  |  |
| 43 | 0.167016 |  |  |  |  |
| 44 | 0.169659 |  |  |  |  |
| 45 | 0.170553 |  |  |  |  |
| 46 | 0.172314 |  |  |  |  |
| 47 | 0.168555 |  |  |  |  |
| 48 | 0.167167 |  |  |  |  |
| 49 | 0.168466 | 0.2 | 0.199 | 0.18 | 0.184 |
| 50 | 0.169854 |  |  |  |  |
| 51 | 0.1698 |  |  |  |  |
| 52 | 0.169595 |  |  |  |  |
| 53 | 0.170667 |  |  |  |  |
| 54 | 0.170184 |  |  |  |  |
| 55 | 0.168718 |  |  |  |  |
| 56 | 0.16588 |  |  |  |  |
| 57 | 0.168414 |  |  |  |  |
| 58 | 0.17022 |  |  |  |  |
| 59 | 0.173457 |  |  |  |  |
| 60 | 0.173288 |  |  |  |  |
| 61 | 0.170883 |  |  |  |  |
| 62 | 0.170718 |  |  |  |  |
| 63 | 0.172162 |  |  |  |  |
| 64 | 0.169682 |  | 0.214 |  | 0.216 |
| 65 | 0.168398 |  |  |  |  |
| 66 | 0.168765 |  |  |  |  |
| 67 | 0.170693 |  |  |  |  |
| 68 | 0.172584 |  |  |  |  |
| 69 | 0.168838 |  |  |  |  |
| 70 | 0.17389 |  |  |  |  |
| 71 | 0.170677 |  |  |  |  |
| 72 | 0.17241 |  |  |  |  |
| 73 | 0.171881 |  |  |  |  |
| 74 | 0.173012 |  |  |  |  |
| 75 | 0.173307 |  |  |  |  |
| 76 | 0.171921 |  |  |  |  |
| 77 | 0.172385 |  |  |  |  |
| 78 | 0.171477 |  |  |  |  |
| 79 | 0.17395 | 0.145 | 0.207 | 0.135 | 0.181 |
| 80 | 0.172792 |  |  |  |  |
| 81 | 0.172502 |  |  |  |  |
| 82 | 0.171482 |  |  |  |  |
| 83 | 0.178029 |  |  |  |  |
| 84 | 0.172957 |  |  |  |  |
| 85 | 0.173607 |  |  |  |  |
| 86 | 0.173609 |  |  |  |  |
| 87 | 0.174501 |  |  |  |  |
| 88 | 0.175742 |  |  |  |  |
| 89 | 0.175057 |  |  |  |  |
| 90 | 0.17235 |  |  |  |  |
| 91 | 0.174769 |  |  |  |  |
| 92 | 0.176034 |  |  |  |  |
| 93 | 0.180073 |  |  |  |  |
| 94 | 0.175832 |  | 0.19 |  | 0.162 |
| 95 | 0.181968 |  |  |  |  |
| 96 | 0.174461 |  |  |  |  |
| 97 | 0.174955 |  |  |  |  |
| 98 | 0.175641 |  |  |  |  |
| 99 | 0.175884 |  |  |  |  |
| 100 | 0.175449 |  |  |  |  |
| 101 | 0.175542 |  |  |  |  |
| 102 | 0.171501 |  |  |  |  |
| 103 | 0.175504 |  |  |  |  |
| 104 | 0.177993 | 0.143 |  | 0.133 |  |
| 105 | 0.18081 |  |  |  |  |
| 106 | 0.179173 |  |  |  |  |
| 107 | 0.182412 |  |  |  |  |
| 108 | 0.183032 |  |  |  |  |
| 109 | 0.180628 |  | 0.178 |  | 0.158 |
| 110 | 0.18241 |  |  |  |  |
| 111 | 0.1818 |  |  |  |  |
| 112 | 0.179725 |  |  |  |  |
| 113 | 0.183959 |  |  |  |  |
| 114 | 0.180352 |  |  |  |  |
| 115 | 0.179855 |  |  |  |  |
| 116 | 0.187742 |  |  |  |  |
| 117 | 0.184972 |  |  |  |  |
| 118 | 0.182812 |  |  |  |  |
| 119 | 0.188231 |  |  |  |  |
| 120 | 0.198713 |  |  |  |  |
| 121 | 0.189132 |  |  |  |  |
| 122 | 0.197247 |  |  |  |  |
| 123 | 0.191015 |  |  |  |  |
| 124 | 0.196831 |  |  |  |  |
| 125 | 0.180199 |  |  |  |  |
| 126 | 0.196019 |  |  |  |  |
| 127 | 0.189387 |  |  |  |  |
| 128 | 0.196187 |  |  |  |  |
| 129 | 0.190886 |  |  |  |  |
| 130 | 0.182067 |  |  |  |  |
| 131 | 0.183573 |  |  |  |  |
| 132 | 0.181514 |  |  |  |  |
| 133 | 0.183561 | 0.12 | 0.162 | 0.115 | 0.143 |

b) Maize- 1

θv(-)

|  | θ(RAW) | M0I1 | M0I2 | M50I1 | M50I2 |
| --- | --- | --- | --- | --- | --- |
| 1 | 0.188588 |  |  |  |  |
| 2 | 0.193029 |  |  |  |  |
| 3 | 0.188598 |  |  |  |  |
| 4 | 0.187903 |  |  |  |  |
| 5 | 0.191435 |  |  |  |  |
| 6 | 0.189177 |  |  |  |  |
| 7 | 0.191298 |  |  |  |  |
| 8 | 0.19268 |  |  |  |  |
| 9 | 0.198769 |  |  |  |  |
| 10 | 0.191423 |  |  |  |  |
| 11 | 0.187525 | 0.2 | 0.202 | 0.22 | 0.213 |
| 12 | 0.184866 |  |  |  |  |
| 13 | 0.185112 |  |  |  |  |
| 14 | 0.188054 |  |  |  |  |
| 15 | 0.19335 |  |  |  |  |
| 16 | 0.193952 |  |  |  |  |
| 17 | 0.187156 |  |  |  |  |
| 18 | 0.186189 |  |  |  |  |
| 19 | 0.178117 |  |  |  |  |
| 20 | 0.185645 |  |  |  |  |
| 21 | 0.186465 |  |  |  |  |
| 22 | 0.188865 |  |  |  |  |
| 23 | 0.184919 |  |  |  |  |
| 24 | 0.185645 |  |  |  |  |
| 25 | 0.186188 |  |  |  |  |
| 26 | 0.187881 | 0.191 | 0.189 | 0.206 | 0.196 |
| 27 | 0.181897 |  |  |  |  |
| 28 | 0.182436 |  |  |  |  |
| 29 | 0.180247 |  |  |  |  |
| 30 | 0.179105 |  |  |  |  |
| 31 | 0.186705 |  |  |  |  |
| 32 | 0.182894 |  |  |  |  |
| 33 | 0.182071 |  |  |  |  |
| 34 | 0.184047 |  |  |  |  |
| 35 | 0.182806 |  |  |  |  |
| 36 | 0.179907 |  |  |  |  |
| 37 | 0.181754 |  |  |  |  |
| 38 | 0.18123 |  |  |  |  |
| 39 | 0.180321 |  |  |  |  |
| 40 | 0.179533 |  |  |  |  |
| 41 | 0.18204 | 0.19 | 0.192 | 0.185 | 0.182 |
| 42 | 0.184145 |  |  |  |  |
| 43 | 0.182133 |  |  |  |  |
| 44 | 0.183041 |  |  |  |  |
| 45 | 0.184368 |  |  |  |  |
| 46 | 0.182929 |  |  |  |  |
| 47 | 0.189047 |  |  |  |  |
| 48 | 0.188754 |  |  |  |  |
| 49 | 0.184266 |  | 0.21 |  | 0.203 |
| 50 | 0.182113 |  |  |  |  |
| 51 | 0.180483 |  |  |  |  |
| 52 | 0.182523 |  |  |  |  |
| 53 | 0.180229 |  |  |  |  |
| 54 | 0.180057 |  |  |  |  |
| 55 | 0.179777 |  |  |  |  |
| 56 | 0.181433 | 0.146 | 0.21 | 0.139 | 0.202 |
| 57 | 0.184991 |  |  |  |  |
| 58 | 0.183041 |  |  |  |  |
| 59 | 0.17716 |  |  |  |  |
| 60 | 0.177158 |  |  |  |  |
| 61 | 0.178605 |  |  |  |  |
| 62 | 0.17917 |  |  |  |  |
| 63 | 0.177632 |  | 0.222 |  | 0.203 |
| 64 | 0.175686 |  |  |  |  |
| 65 | 0.173367 |  |  |  |  |
| 66 | 0.172106 |  |  |  |  |
| 67 | 0.172456 |  |  |  |  |
| 68 | 0.172616 |  |  |  |  |
| 69 | 0.172654 |  |  |  |  |
| 70 | 0.171946 |  |  |  |  |
| 71 | 0.179624 | 0.151 |  | 0.148 |  |
| 72 | 0.171406 |  |  |  |  |
| 73 | 0.173113 |  |  |  |  |
| 74 | 0.172925 |  |  |  |  |
| 75 | 0.174116 |  |  |  |  |
| 76 | 0.173188 |  |  |  |  |
| 77 | 0.174136 |  |  |  |  |
| 78 | 0.177824 |  |  |  |  |
| 79 | 0.170658 |  | 0.196 |  | 0.19 |
| 80 | 0.168967 |  |  |  |  |
| 81 | 0.169707 |  |  |  |  |
| 82 | 0.170403 |  |  |  |  |
| 83 | 0.168868 |  |  |  |  |
| 84 | 0.169443 |  |  |  |  |
| 85 | 0.169712 |  |  |  |  |
| 86 | 0.176059 |  |  |  |  |
| 87 | 0.170704 |  |  |  |  |
| 88 | 0.168481 |  |  |  |  |
| 89 | 0.167511 |  |  |  |  |
| 90 | 0.167926 |  |  |  |  |
| 91 | 0.167553 |  |  |  |  |
| 92 | 0.166166 |  |  |  |  |
| 93 | 0.167127 |  |  |  |  |
| 94 | 0.169751 | 0.15 | 0.24 | 0.141 | 0.223 |
| 95 | 0.168391 |  |  |  |  |
| 96 | 0.16672 |  |  |  |  |
| 97 | 0.167494 |  |  |  |  |
| 98 | 0.166229 |  |  |  |  |
| 99 | 0.168193 |  |  |  |  |
| 100 | 0.165062 |  |  |  |  |
| 101 | 0.166114 |  |  |  |  |
| 102 | 0.171021 |  |  |  |  |
| 103 | 0.17001 |  |  |  |  |
| 104 | 0.166083 |  |  |  |  |
| 105 | 0.166271 |  |  |  |  |
| 106 | 0.165661 |  |  |  |  |
| 107 | 0.165283 |  |  |  |  |
| 108 | 0.165138 |  |  |  |  |
| 109 | 0.165945 |  |  |  |  |
| 110 | 0.169313 |  |  |  |  |
| 111 | 0.171188 |  |  |  |  |
| 112 | 0.166036 |  |  |  |  |
| 113 | 0.168397 |  |  |  |  |
| 114 | 0.168025 |  |  |  |  |
| 115 | 0.168137 | 0.24 | 0.265 | 0.235 | 0.255 |

d) Maize-2

|  | θ(RAW) | M0I1 | M0I2 | M50I1 | M50I2 |
| --- | --- | --- | --- | --- | --- |
| 1 | 0.180706 |  |  |  |  |
| 2 | 0.178817 |  |  |  |  |
| 3 | 0.179524 |  |  |  |  |
| 4 | 0.188096 |  |  |  |  |
| 5 | 0.183967 |  |  |  |  |
| 6 | 0.186754 |  |  |  |  |
| 7 | 0.183838 |  |  |  |  |
| 8 | 0.185073 |  |  |  |  |
| 9 | 0.188915 |  |  |  |  |
| 10 | 0.184212 |  |  |  |  |
| 11 | 0.188196 | 0.19 | 0.203 | 0.21 | 0.211 |
| 12 | 0.185232 |  |  |  |  |
| 13 | 0.185975 |  |  |  |  |
| 14 | 0.190387 |  |  |  |  |
| 15 | 0.188645 |  |  |  |  |
| 16 | 0.194017 |  |  |  |  |
| 17 | 0.185149 |  |  |  |  |
| 18 | 0.191322 |  |  |  |  |
| 19 | 0.188762 |  |  |  |  |
| 20 | 0.18858 |  |  |  |  |
| 21 | 0.188996 |  |  |  |  |
| 22 | 0.183103 |  |  |  |  |
| 23 | 0.18753 |  |  |  |  |
| 24 | 0.188731 |  |  |  |  |
| 25 | 0.194904 |  |  |  |  |
| 26 | 0.187189 | 0.188 | 0.183 | 0.193 | 0.195 |
| 27 | 0.187047 |  |  |  |  |
| 28 | 0.180394 |  |  |  |  |
| 29 | 0.190115 |  |  |  |  |
| 30 | 0.191523 |  |  |  |  |
| 31 | 0.191774 |  |  |  |  |
| 32 | 0.182606 |  |  |  |  |
| 33 | 0.181967 |  |  |  |  |
| 34 | 0.173354 |  |  |  |  |
| 35 | 0.175698 |  |  |  |  |
| 36 | 0.176538 |  |  |  |  |
| 37 | 0.177067 |  |  |  |  |
| 38 | 0.178054 |  |  |  |  |
| 39 | 0.187334 |  |  |  |  |
| 40 | 0.184023 |  |  |  |  |
| 41 | 0.179646 | 0.172 | 0.174 | 0.184 | 0.194 |
| 42 | 0.178533 |  |  |  |  |
| 43 | 0.179315 |  |  |  |  |
| 44 | 0.177565 |  |  |  |  |
| 45 | 0.1801 |  |  |  |  |
| 46 | 0.177401 |  |  |  |  |
| 47 | 0.177302 |  |  |  |  |
| 48 | 0.178206 |  |  |  |  |
| 49 | 0.179359 |  | 0.213 |  | 0.203 |
| 50 | 0.175843 |  |  |  |  |
| 51 | 0.176742 |  |  |  |  |
| 52 | 0.175223 |  |  |  |  |
| 53 | 0.173621 |  |  |  |  |
| 54 | 0.175843 |  |  |  |  |
| 55 | 0.174707 |  |  |  |  |
| 56 | 0.173524 | 0.153 | 0.21 | 0.136 | 0.196 |
| 57 | 0.173608 |  |  |  |  |
| 58 | 0.173356 |  |  |  |  |
| 59 | 0.173362 |  |  |  |  |
| 60 | 0.175192 |  |  |  |  |
| 61 | 0.175076 |  |  |  |  |
| 62 | 0.174444 |  |  |  |  |
| 63 | 0.169723 |  | 0.22 |  | 0.196 |
| 64 | 0.169248 |  |  |  |  |
| 65 | 0.168339 |  |  |  |  |
| 66 | 0.169655 |  |  |  |  |
| 67 | 0.169331 |  |  |  |  |
| 68 | 0.169916 |  |  |  |  |
| 69 | 0.166742 |  |  |  |  |
| 70 | 0.167745 |  |  |  |  |
| 71 | 0.169951 | 0.147 |  | 0.136 |  |
| 72 | 0.17201 |  |  |  |  |
| 73 | 0.171013 |  |  |  |  |
| 74 | 0.166519 |  |  |  |  |
| 75 | 0.168202 |  |  |  |  |
| 76 | 0.167616 |  |  |  |  |
| 77 | 0.167396 |  |  |  |  |
| 78 | 0.166913 |  |  |  |  |
| 79 | 0.16861 |  | 0.201 |  | 0.179 |
| 80 | 0.17283 |  |  |  |  |
| 81 | 0.170864 |  |  |  |  |
| 82 | 0.166987 |  |  |  |  |
| 83 | 0.166873 |  |  |  |  |
| 84 | 0.166277 |  |  |  |  |
| 85 | 0.168246 |  |  |  |  |
| 86 | 0.170508 |  |  |  |  |
| 87 | 0.170672 |  |  |  |  |
| 88 | 0.171277 |  |  |  |  |
| 89 | 0.166101 |  |  |  |  |
| 90 | 0.16882 |  |  |  |  |
| 91 | 0.168895 |  |  |  |  |
| 92 | 0.167567 |  |  |  |  |
| 93 | 0.167809 |  |  |  |  |
| 94 | 0.166877 | 0.149 | 0.223 | 0.147 | 0.197 |
| 95 | 0.167966 |  |  |  |  |
| 96 | 0.168005 |  |  |  |  |
| 97 | 0.168435 |  |  |  |  |
| 98 | 0.167422 |  |  |  |  |
| 99 | 0.165537 |  |  |  |  |
| 100 | 0.165081 |  |  |  |  |
| 101 | 0.167028 |  |  |  |  |
| 102 | 0.164316 |  |  |  |  |
| 103 | 0.163226 |  |  |  |  |
| 104 | 0.167603 |  |  |  |  |
| 105 | 0.165752 |  |  |  |  |
| 106 | 0.164187 |  |  |  |  |
| 107 | 0.167422 |  |  |  |  |
| 108 | 0.165537 |  |  |  |  |
| 109 | 0.165081 |  |  |  |  |
| 110 | 0.167028 |  |  |  |  |
| 111 | 0.164316 |  |  |  |  |
| 112 | 0.163226 | 0.2459 | 0.263 | 0.267 | 0.267 |

**Figure S2.** Moisture measured before each irrigation, and critical limit of readily available water.

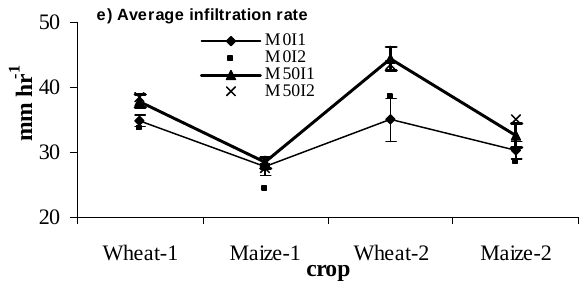

**Figure S3.** Effect of manure and irrigation levels on soil saturated hydraulic conductivity at harvest of wheat and maize Note: Year effect was statistically non-significant (*T*-test) at harvest of wheat and maize.
